# A multifaceted microRNA turnover complex from *Caenorhabditis elegans*

**DOI:** 10.1101/2020.12.28.424526

**Authors:** Mohini Singh, Adil R. Wani, Pradipta Kundu, Saibal Chatterjee

## Abstract

microRNAs are known to regulate expression of more than two third of all the eukaryotic genes by post-transcriptional means, and regulation of these tiny regulators play an important role in determining their activities. Here, we report a macromolecular microRNA turnover complex, whose components are crucial to microRNA homeostasis and development in *Caenorhabditis elegans*. Biochemical investigations with the purified complex in an isolated system not only unfolded the roles of the individual subunits critical for the functionality of the complex, but also unraveled the different modes of operations and regulatability of this biological machine. Our results reveal that this complex is highly receptive and capable of switching between an ATP-dependent and ATP-independent mode of operation depending on the availability of ATP in its environment, which might allow the complex to function dynamically during different physiological conditions.

---

microRNAs (miRNAs) are endogenous small non-coding RNAs, which constitute a major layer of eukaryotic gene regulation by modulating post-transcriptional expression of a large number of mRNAs through employing antisense mechanisms, and thus collectively regulate diverse developmental and physiological processes (*1*). Recent reports have indicated that differential turnover plays a crucial role in determining their abundance, and thus functionality of miRNAs (*1, 2*). Although, biogenesis of miRNAs has been extensively studied (*1, 3*), but very little is known about the basic miRNA turnover pathway, let alone how its differential regulation can bring about different physiological outcomes.

The first report on miRNA turnover was based on a work involving the plant system *Arabidopsis thaliana*, where Small RNA Degrading Nuclease1 (SDN1) was shown to degrade a handful of miRNAs (*4*). Then it was reported that miRNA turnover is an active process in *Caenorhabditis elegans* (*C. elegans*), where 5’-3’ exoribonuclease-2 (XRN-2) acts as a ‘miRNase’ (*5*). Later partner of XRN two-1 (PAXT-1) was shown as its co-factor (*6*), which contributes by stabilizing the former. XRN-1 was also found to play the role of a ‘miRNase’ in the worms (*7*), and subsequently decapping scavenger enzyme 1 (DCS-1) was shown to stimulate the ‘miRNase’ activity of XRN-1 (*8*).

miRNA turnover pathway in the human cells is also not thoroughly understood. Initial studies using HeLa cells reported varying stability of miR-29b during different cell cycle states (*9*). Intrinsic sequences were also found to facilitate or promote the decay of a given miRNA (*9-11*). It was reported that the exosome complex and XRN1 mediate decay of miR-382 in HEK293 cells (*10*), whereas PNPase old-35 mediates the decay of miR-221, miR-222, miR-106b in HO-1 cells (*12*). In HEK 293T cells Tudor-SN was shown to be the ‘miRNase’ for a cohort of miRNAs, which play a crucial role in G1 to S phase transition (*13*). Recently an unidentified 3’ trimming activity has been implicated in target RNA directed degradation of miR-29b in vertebrate cerebellum (*14*). But, all these reports still do not paint a coherent picture of the core pathway, the cascade of interactions, and the operational mechanisms employed by the ‘units’ of the miRNA turnover pathway in animals.

Notably, the studies in *C. elegans* made some significant advancement by showing how miRNA turnover is a stepwise process, where as-yet-unknown activity releases the miRNA, before its degradation by XRN-2 (*5*). It was also demonstrated that miRNA targets can modulate the degradation of a miRNA (*5, 7*), and furthermore, how it might contribute to the process of evolution of miRNAs (*7*). But, both the ‘miRNases’ identified in *C. elegans* have also been implicated in the processing and turnover of a wide variety of RNAs across evolution (*2, 15, 16*). As a result, our understanding of the regulation of the ‘miRNase’ activity of XRN-1 and XRN-2 remained unclear and illusive, and at the same time their potential to be used as therapeutic targets to control deregulated miRNA levels in different disease-states appeared much limited (*2, 17*). Many of the eukaryotic nucleases, and almost all of the nucleases playing crucial roles in the small RNA pathways have been found to be residing in large complexes (*1, 3, 18*), associated with defined sets of co-factors, which direct them to specific pathways, determine their specificities, and achieve responsiveness to regulatory cues to bring about different physiological outcomes. Thus, it is possible that ‘miRNases’ also reside in macromolecular complexes, where the molecular niche partners govern their roles by conferring specificity and regulatability. And this possibility was bolstered by the observation that endogenous XRN-1/2 in *C. elegans* predominantly reside in high molecular weight complexes (see results) rather than as ‘individual species’. Therefore, purification followed by unraveling of the mode of action of the miRNA turnover complexes would be crucial to understand miRNA turnover pathway, rather than studying the individual ‘miRNases’ in isolation.

In this study, we report a macromolecular miRNA turnover complex, miRNasome-1, whose components are crucial to miRNA homeostasis and development in *C. elegans*. The complex is composed of four subunits including XRN-2-the ‘miRNase’, and here we also report a previously unknown endoribonuclease activity of this fundamentally important enzyme. Our results show that miRNasome-1-residing XRN-2’s activity and specificity is governed by two of the newly identified members of this biological machine. The RNA-binding receptor component of the complex is not only found to be crucial for worm development, but it also confers *in vivo* substrate specificity to the complex, which corroborates with that of the complex’s activity, *in vitro*. And an ATP binding/ hydrolysing subunit helps the complex to switch between two alternative mechanisms of turnover (energy-independent endoribonucleolysis vs energy-dependent exoribonucleolysis). Biochemical assays with the purified complex in an isolated system facilitated the unraveling of its receptive *modi operandi* and regulatability.

## RESULTS

### Purification of miRNasome-1 and its activity

In order to elucidate the modus operandi of a ‘miRNase’, in its actual molecular niche, which would essentially form a functional unit of the miRNA turnover pathway and an integral part of the miRNA metabolism circuitry in *C. elegans*, we decided to purify the ‘miRNases’ from endogenous sources along with the members of their molecular niche. From the biochemical properties of miRNA turnover *in vitro* in *C. elegans*, we identified a potential tool for their purification, and further incorporated that into an effective purification strategy. We had observed that in an *in vitro* miRNA turnover assay (*5*), RNAs slightly shorter or longer than the miRNA size-range are relatively resistant to total worm lysate (**Fig. 1A**). That indicated to the existence of some machinery selecting its substrate by size, which was reminiscent of the well-known viral suppressor proteins recognizing host-siRNAs by size (*19, 20*). This observation suggested that the mature miRNA, if stabilized, might act as an affinity arm for purification of the activity acting on miRNAs. Indeed, a phosphorothioate (PTO) stabilized miRNA (*let-7*) could be utilized as an affinity arm to purify a high molecular weight protein complex (∼260 kDa, **fig. S1A, B**), which showed specific activity on miRNA (**fig. S1C-E**), albeit the yield was poor, and was not homogeneous (**fig. S1B top panel**). Western blotting revealed the presence of the known ‘miRNase’-XRN-2 in the purified complex, but XRN-1 appeared to be absent (compare **fig. S1B middle and bottom panels**). In order to improve the purification, we introduced a fractionation step (ion-exchange, **Fig. 1B**, and **fig. S2**) prior to RNA-affinity purification (**Fig. 1C**). This particular combination of chromatographic methods resulted in the purification of an XRN-2 containing high molecular weight band to purity (**Fig. 1B, C;** see methods for details). Our starting material was total lysate from the worm larval stage 4 (L4), and this was aimed to capture the miRNA turnover complex, which might be playing a crucial role in shaping the miRNA landscape facilitating the larval to adult transition. Importantly, mammalian homologues of many of those *C. elegans* L4 stage miRNAs are involved in critical cellular and developmental processes (eg. *let-7* in cellular proliferation and differentiation, references *21-23*), whose deregulation leads to several disease states including cancer. The purified macromolecular protein complex was composed of four subunits in the apparent stoichiometry of 1:1:1:1 for each of the four (**Fig. 1D**), and with an aggregate molecular mass of ∼261 kDa. We christened this miRNA turnover complex “miRNasome-1”, since it is the first such endogenous protein complex to be purified.

**Fig 1.**
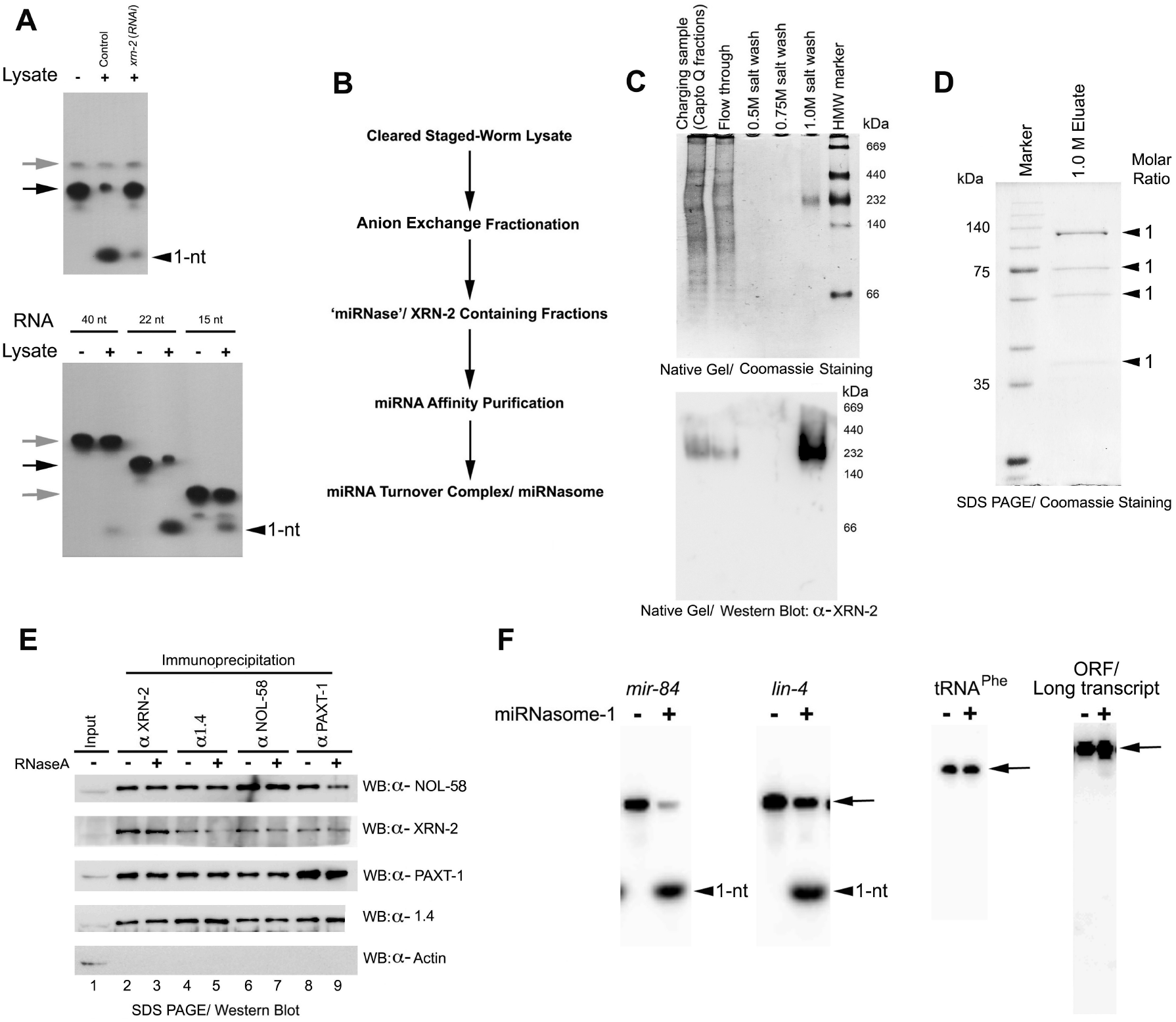
Purification of miRNasome-1 and its activity. **(A) XRN-2 dependent miRNA turnover activity in worm lysate governed by RNA size**. (**Top panel**) Efficient turnover of 3’-pCp-labeled and blocked, and 5’-monophosphorylated mature *let-7* substrate (black arrow) in the control lysate, when product (arrowhead) formation got heavily diminished in XRN-2 depleted (RNAi) lysate. Grey arrowhead indicates to the larger artifactual RNA band, which remained largely unaffected. (**Bottom panel**) 5’-monophosphorylated RNAs of the indicated sizes, similarly radiolabeled as above, were subjected to turnover assay. The 22-nucleotide (nt) long RNA (*let-7*, black arrow) gets efficiently converted into product (arrowhead), whereas the longer and shorter substrates (grey arrows) produce very little product (arrowhead). **(B)** Purification scheme comprising of ion exchange and RNA-affinity chromatography, where stabilized miRNA (*let-7*) was used as an affinity-arm. **(C)** The fractions of interest from the Capto Q purification step were subjected to RNA affinity chromatography, and the high-salt (1.0 M) eluate resolved on a native protein gel shows a single high molecular weight band (miRNasome-1, **top panel**). Western blot confirmed the band to harbor the ‘miRNase’-XRN-2 (**bottom panel**). **(D)** SDS PAGE analysis of the RNA affinity chromatography purified miRNasome-1 reveals the presence of four subunits in the apparent stoichiometry of 1:1:1:1. Note that the presence of salt in the 1.0 M eluate resulted in slower than expected migration for the proteins. **(E)** Immunoprecipitation (IP) of endogenous proteins, as indicated, followed by their western blot analyses using antibodies against XRN-2, PAXT-1, NOL-58, and miRNasome-1.4. XRN-2 is detected in PAXT-1, NOL-58 and miRNasome-1.4 immunoprecipitates and vice versa. RNase A treatment does not perturb the interactions between the subunits. 1% of input and 100% of the eluates were subjected to western analyses. In this figure and the subsequent concerned figures, α-miRNasome-1.4 antibody has been represented as α-1.4. Note that the differences in migration between input and IP samples are caused by the differences in the salt content. **(F)** Activity of purified miRNasome-1 (100 ng, ∼40 nM) on different 5’-radiolabeled mature miRNAs, tRNA, and body-radiolabeled *Renilla* luciferase ORF (∼1100-nt), as indicated. Substrate RNA concentration in all the above reactions was 200 nM. Substrates are indicated with arrows, and products with arrowheads. Representative images from three or more repeats have been presented.

Mass spectrometric analysis of the purified complex identified four proteins from the excised single-band, obtained upon resolving the purified sample on a native gel (**Table S1**). Amongst them, two were already reported factors involved in miRNA turnover (XRN-2 and PAXT-1, references *5, 6*), and out of the two newly identified proteins one (NOL-58) is an ortholog of human NOP58, which is a structural subunit of the C/D Box snoRNPs (*24*). The other one (B0024.11) is predicted to be an ortholog of human PUS7 (*24*), which is a putative pseudouridylate synthase (henceforth, named as miRNasome-1.4). These two proteins get expressed ubiquitously during the worm life cycle, but their detailed functions and the mechanisms employed by them are not well understood (*24*). The interaction between XRN-2, PAXT-1 and these newly identified partners could also be suggested from co-immunoprecipitation experiments using total worm lysate, and resistance to RNase A-treatment confirmed their direct physical interaction (**Fig. 1E**, see materials & methods). Moreover, the recombinant versions of the miRNasome-1 subunits could reconstitute a macromolecular complex, where it could be suggested that the individual subunits were present in the apparent stoichiometry of 1:1:1:1 (**fig. S3)**.

In *in vitro* turnover assays, endogenous miRNAsome-1 showed robust activity on specific miRNAs, and very little or no activity on other types of RNAs (tRNA, ORF/ long transcript, **Fig. 1F**). These observations were in contrast to the activity of recombinant XRN-2, which acted efficaciously on miRNA and ORF/ long transcript, but showed negligible activity on tRNA, in *in vitro* assays (**fig. S4A-C**). Notably, in expected lines, the *in vitro* reconstituted complex also showed substrate specificity similar to that of the endogenous miRNasome-1 (**fig. S4D-F**).

### *nol-58*(*RNAi*) suppresses *let-7* mutant phenotype

To unravel the physiological significance of the newly identified interacting-partners of the ‘miRNase’, we resorted to the worm strain *let-7*(*n2853*) (reference *25*), which has a single point mutation in the seed sequence of *let-7*, additionally the level of mature *let-7* is 2-3 fold diminished compared to that of the wild type (*5, 25*). All these result into greater abundance of *let-7* targets, and the worm dies by bursting through its vulva during the L4 to Adult transition, when grown at the non-permissive temperature (25^°^C, references *5, 25*). We hypothesized that if the newly identified factors would play a direct role in miRNA turnover, then their depletion would increase miRNA levels, including *let-7*, and that might bring down the levels of the deregulated targets to alleviate the bursting phenotype. Indeed, RNAi depletion of NOL-58 not only led to the efficient suppression (79% **±** 2.5% [n=200]) of the bursting phenotype (**fig. S5**, references *5, 24*), but also allowed the generation of adult alae (73% **±** 4.0% [n=50]); **fig. S5A** and **B** bottom panels), a cuticular structure, whose formation is dependent on *let-7* functioning (**Supplementary Text 1, fig. S6**, reference *25*). Of note, *nol-58* is already reported to be a suppressor of *let-7*(*n2853*) lethality (*26*).

Employing additional *in vivo* tools (*6, 27*), *nol-58* could also be shown to be a genetic enhancer of *xrn-2* (**Supplementary Text 2, fig. S7**). However, *miRNasome-1*.*4* didn’t prove to be a genetic enhancer of *xrn-2*, nor did its depletion alleviate the bursting phenotype of the *let-7*(*n2853*) worms (data not shown). And that could be due to its possible accessory role or a regulatory role meant for a given physiological situation.

### NOL-58 affects functional mature miRNAs *in vivo*

To gain molecular evidence for a role of *let-7* in the *let-7*(*n2853*) worms undergoing *nol-58*(*RNAi*), we extracted total RNA from L4 stage worms, alongside control worms of the exact same stage, and subjected them to northern blotting using anti-*let-7* probe. There was ∼3 fold accumulation of mature *let-7*, and without any change in the levels of the pre-*let-7*, as well as pri-*let-7* (**Fig. 2A, B**). More importantly, when we measured the levels of two of the validated *let-7* targets (*5, 25, 28, 29*) through RT-qPCR, we observed that those targets in the NOL-58 depleted worms have dropped dramatically compared to that of the control *let-7*(*n2853*) worms. Target levels in those experimental samples were comparable to that of the wild type *N2* worms (**Fig. 2C**). Additionally, levels of control non-target mRNAs (eg. *act-1, tbb-1*) didn’t change upon *nol-58*(*RNAi*) (data not shown), which excluded the possibility that NOL-58 depletion exerted some indirect effect on the levels of the target mRNAs (eg. by affecting transcription), alongside miRNA-mediated direct regulation. Thus, it clearly indicated that the observed accumulated mature *let-7* was functional, and NOL-58 is part of an active turnover mechanism, which affects active mature form of the *let-7* and not its precursors. Additionally, northern blotting also revealed the accumulation of several other miRNAs beyond *let-7*, and without any effect on the levels of their precursors, wherever we could detect them (**Fig. 2A, B**). Similar effects, only on the levels of mature miRNAs, but not all miRNAs, were also observed through northern blotting upon depletion of NOL-58 in the wild type *N2* worms (**fig. S8**). The effects of NOL-58 depletion on mature miRNA levels could also be reconfirmed through independent methods like TaqMan assays (data not shown). Consistent with previous observations (*5*), *xrn-2*(*RNAi*) also affected mature miRNAs only (**Fig. 2A, fig. S8**). Conversely, depletion of miRNasome-1.4 didn’t show any significant effect on miRNA levels (**fig. S8**). Of note, RNAi depletion of NOL-58 didn’t exert any significant off-target effect on the endogenous levels of other miRNasome-1 components, as well as the other known ‘miRNase’-XRN-1, and miRNA-Argonautes (ALG-1, ALG-2), which essentially confirmed the aforementioned effects to be specific (**fig. S9**).

**Fig 2.**
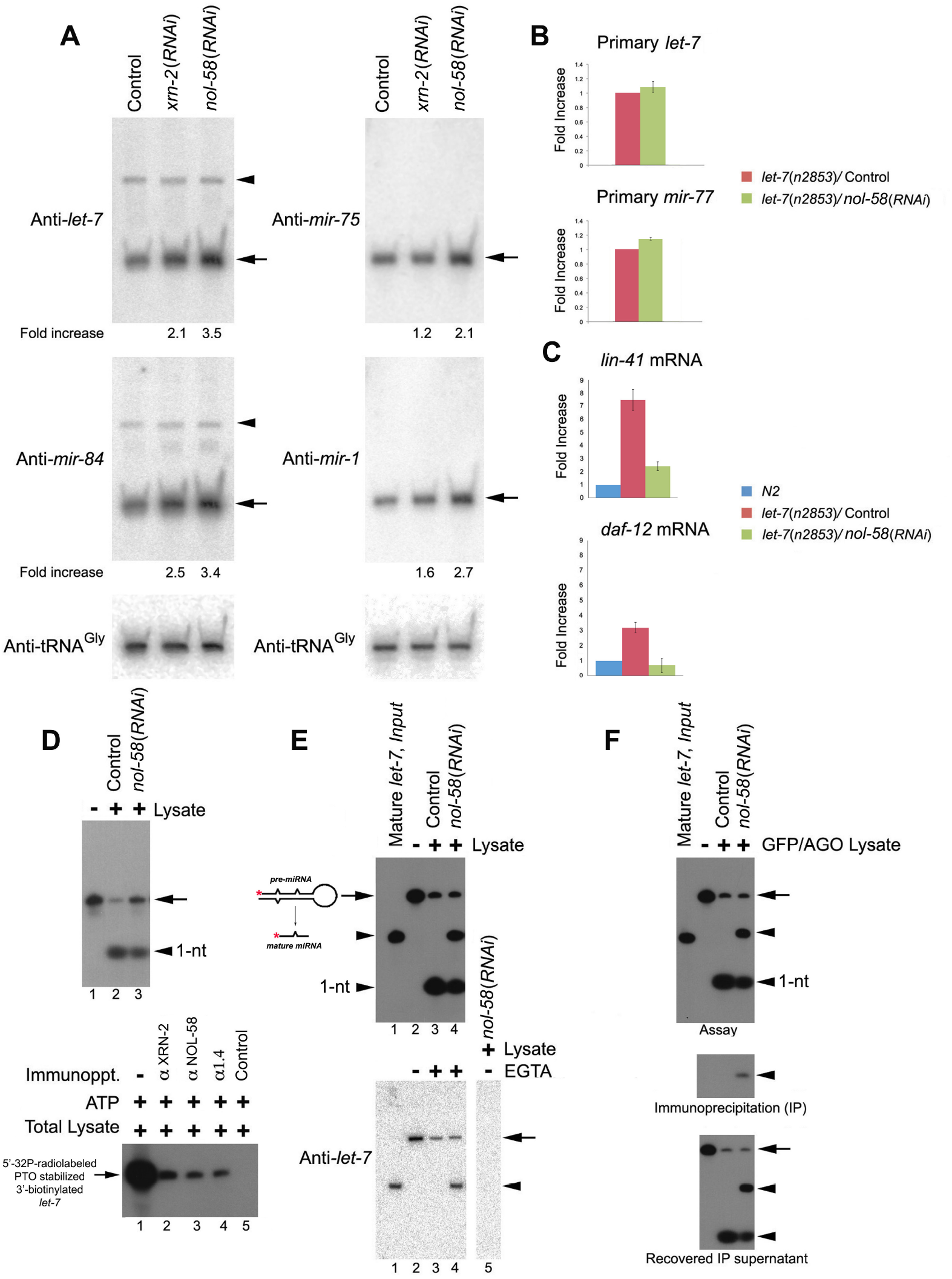
NOL-58 affects functional mature miRNAs *in vivo* and it coordinates with the miRNA precursor-processing machinery *in vitro*. **(A)** RNAi depletion of indicated miRNasome-1 components, as revealed through northern blotting, leads to the accumulation of mature miRNAs (arrow), without affecting the pre-miRNAs (arrowhead). The fold increase in mature miRNA levels has been indicated below the relevant blots. tRNA^Gly^ served as loading control. **(B)** RT–qPCR (n = 3; means ± SEM) ascertains no significant change in the levels of pri-miRNAs from samples, as indicated. **(C)** Determination of the levels of *let-7* targets, *lin-41* and *daf-12*, using RT–qPCR (n = 3; means ± SEM) from samples, as indicated. **(D) Top panel**. NOL-58 affects miRNA stability *in vitro*. Incubation of 3’-pCp-labeled and blocked synthetic mature *let-7* (arrow) with worm lysates, as indicated, largely yields a single product (monoribonucleotide, arrowhead). **Bottom panel**. Single stranded radiolabeled *let-7* miRNA directly interacts with miRNasome-1 components. Appropriately modified *let-7* miRNA was incubated with total worm lysate under cold conditions and subjected to immunoprecipitation using antibodies against miRNasome-1 subunits, as indicated. RNA was extracted from all the immunoprecipitates and subjected to gel electrophoresis. Lane 1 depicts the input miRNA, which was incubated with the total lysate and directly subjected to RNA extraction without proceeding to immunoprecipitation. A representative image from three repeats has been presented. **(E) Top panel**. Coupled pre-*let-7* processing and mature miRNA turnover. 5’-radiolabeled synthetic pre-*let-7* (mature *let-7* resides in the 5’-end of this precursor) was incubated with lysates as indicated. RNAi depletion of NOL-58 leads to the accumulation of mature miRNA (upper arrowhead). The 5’-PNK-radiolabeled (red asterisk) pre-*let-7* and the *in vitro* diced/ processed mature *let-7* from the precursor are depicted as a cartoon on the left. **Bottom panel**. 5’-monophosphorylated cold pre-*let-7* was incubated with the lysates, as indicated, and subjected to northern probing using anti-*let-7* probe. All lanes are from the same blot. **(F) top panel**. pre-*let-7* assay, as in (**E**), performed with GFP/AGO worms undergoing RNAi, as indicated. **middle panel**. *in vitro* processed mature *let-7* co-immunoprecipitates with GFP/AGO. **bottom panel**. Recovered supernatant after immunoprecipitation. All lanes are from the same gel. Representative images from three or more repeats have been presented.

### NOL-58 coordinates with miRNA biogenesis machinery *in vitro*

Towards understanding the role of *nol-58* biochemically, we performed *in vitro* miRNA turnover assays (*5*) using 3’-pCp radiolabeled-and-blocked, and 5’-monophosphorylated synthetic *let-7* and lysate from worms undergoing mock or RNAi for the candidate gene. As reported before (*5, 8*), most of the miRNA got converted into exoribonucleolytic product, monoribonucleotides, in the control lysate (**Fig. 2D top panel**), without the formation of any major intermediates. Depletion of NOL-58 diminished the formation of monoribonucleotides, leaving a considerable amount of intact *let-7*, thus indicating that NOL-58, like XRN-2 (*5*), also exerts an effect on miRNA stability *in vitro* (**Fig. 2D top panel, compare lanes 2, 3**). Moreover, we could co-immunoprecipitate a 5’-radiolabeled and PTO stabilized *let-7* using antibodies against NOL-58 and other miRNasome-1 subunits, where the labeled miRNA was subjected to incubation with total lysate, followed by immunoprecipitation. That confirmed a direct interaction between single stranded mature miRNA and miRNasome-1 (**Fig. 2D bottom panel**). However, since, mature miRNAs are known to be generated from pre-miRNAs in tightly coupled reactions, we decided to perform *in vitro* pre-miRNA processing assays (*5*) using a 5’-radiolabeled synthetic pre-*let-7* and control or experimental worm lysates, and follow the fate of the *in vitro* diced mature miRNA, which would help us to better assess the role of a candidate gene. Incubation with the control lysate led to the disappearance of most of the substrate pre*-let-7* (**Fig. 2E top panel**), and concomitant appearance of a product (**Fig. 2E top panel**), which could be identified as monoribonucleotide through TLC analysis (data not shown). We observed stabilization of radiolabeled pre-*let-7*, when it was incubated with a *dcr-1*(*RNAi*) lysate, which indicated that the product formation in the control lysate was dependent on Dicer cleavage (**fig. S10**). When NOL-58 depleted lysate was used, although the substrate band still disappeared, but there was diminished monoribonucleotide formation and a concomitant appearance of a new band, which co-migrated with a 5’-radiolabeled synthetic mature *let-7* (**Fig. 2E top panel, compare lanes 3, 4**). An independent experiment identified this band as mature *let-7* (**Fig. 2E bottom panel, Supplementary Text 3**).

Since, it is already known that Dicer processing and RISC loading is a pre-requisite for mature miRNA turnover in pre-miRNA processing assays (*5*), the above results indicated that NOL-58 is part of the *bona fide* miRNA turnover machinery, which coordinates with upstream machinery hosting the mature miRNA after pre-miRNA processing, i.e.; the miRISC complex. And this notion was indeed confirmed, when we could show the association of miRNA-argonautes and the *in vitro* processed mature *let-7*, which accumulated in the pre-*let-7* processing assay performed with the lysate depleted for NOL-58 (**Fig. 2F**). In this assay we used worm strains (*30*), where miRNA-Argonautes, ALG-1 and ALG-2, are tagged with GFP (GFP/AGO), which allows immunoprecipitation of the GFP/AGO using anti-GFP antibody and analysis of their RNA content (*5*). A modest amount of *in vitro* processed 5’-radiolabeled mature *let-7* could be co-immunoprecipitated using anti-GFP antibody (**Fig. 2F middle panel**), and the majority of the mature *let-7* was found in the sup-fraction (**Fig. 2F bottom panel**). Such low miRNA recovery could well be due to the limit of the immunoprecipitation efficiency (∼10-15% of the input). Additionally, depletion of NOL-58 significantly increased the amount of endogenous mature miRNAs co-immunoprecipitated with GFP/AGO, which again suggested that miRNA release from miRNA-Argonautes, and its subsequent turnover by miRNasome-1 are kinetically linked processes (data not shown).

### An ATP-independent endoribonuclease activity of miRNasome-1

In order to decipher the modus operandi, and to understand the regulatability of miRNasome-1, we performed several *in vitro* biochemical assays using radiolabeled miRNA and purified endogenous miRNasome-1. Here we used a twenty fold lower concentration of protein than that used before in similar assays (**Fig. 1F**), so that the intermediates could be detected, if any. Upon incubation with a five-fold molar excess of a body-radiolabeled substrate (*mir-84*), in the absence of ATP, a single product/ band was formed, which was found to be 3-nucleotide (nt) long (**Fig. 3A top panel** and **3B top left panel**). This product formation was gradually inhibited with progressive increase in ATP in the reaction (**Fig. 3A top panel**), but without any significant formation of monoribonucleotides.

**Fig 3.**
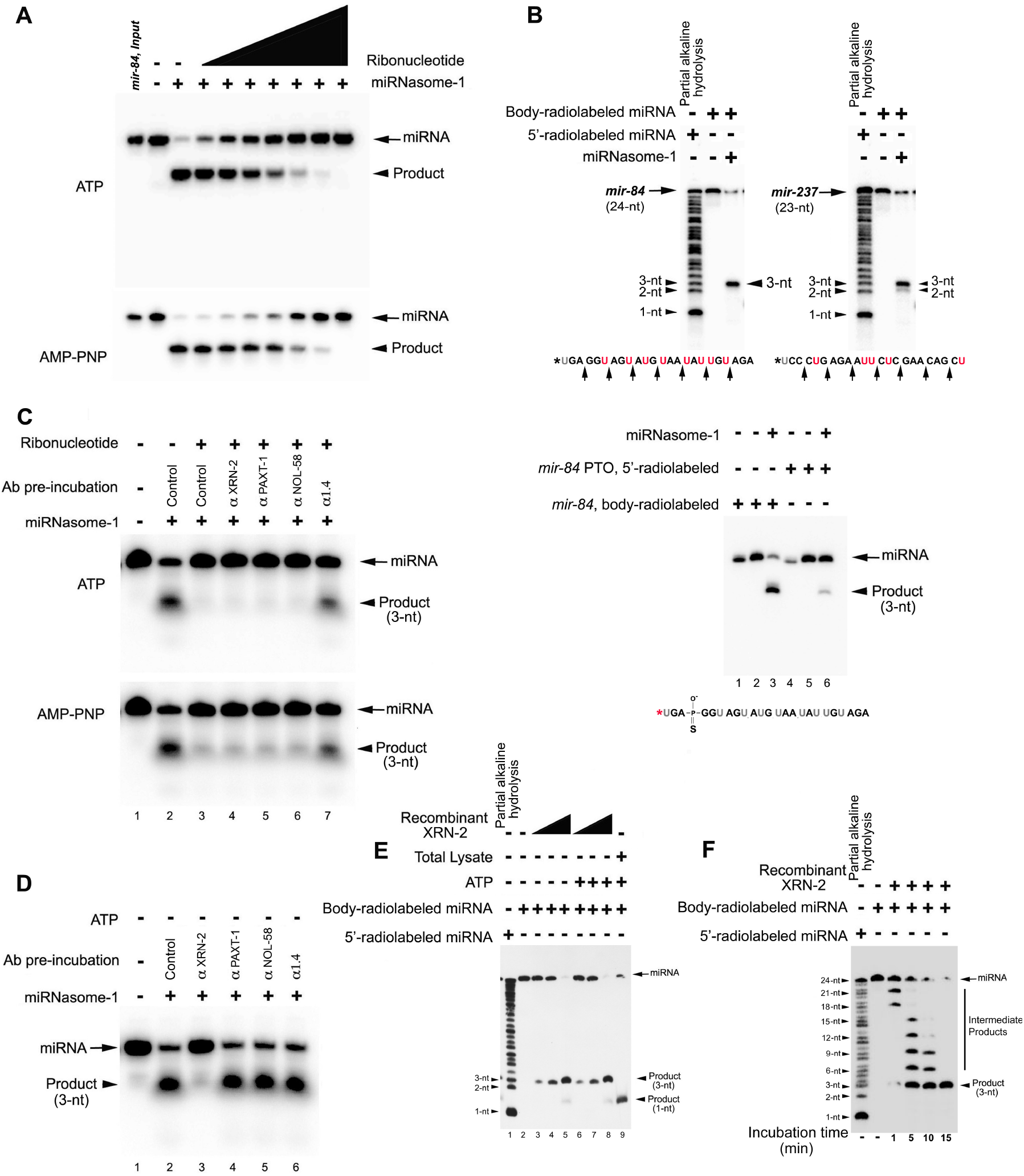
XRN-2 confers an ATP-independent endoribonucleolytic activity to miRNasome-1. **(A)** Inhibitory effect of ribonucleotides on product formation by miRNasome-1. Body-labeled *mir-84* (100 fmol,10 nM) subjected to *in vitro* miRNA turnover assay performed with purified miRNasome-1 (5 ng, ∼1.9 nM) in the absence or presence of an ascending concentration (0.25 mM - 3 mM) of ATP (**top panel**) or AMP-PNP (**bottom panel**), as indicated. **(B) Top panel**. miRNasome-1 performs highly processive endoribonucleolytic cleavage after every 3-nt in the absence of ATP. miRNasome-1 was incubated with a body-radiolabeled and 5’-phosphorylated *mir-84* and *mir-237*(arrow) in the absence of ATP, and resolved on urea PAGE, alongside partial alkaline hydrolysate of respective 5’-radiolabeled miRNAs, as indicated. The relevant miRNA sequence and the putative endoribonucleolytic cleavage sites (indicated by arrows) are furnished below each gel picture. Asterisk represents 5’-phosphate (cold) end of a miRNA. **Bottom panel**. Stabilization of an internal bond between the 3^rd^ and 4^th^ nucleoside of *mir-84* diminishes the product formation by miRNasome-1 in the absence of ATP. miRNAs, as indicated, were incubated with miRNasome-1 in the absence of ATP, and the reaction products were resolved on a urea PAGE. Same molar concentration was used for both the miRNAs. **(C)** Antibody against miRNasome-1.4 prevents inhibitory effect of ribonucleotides on the endoribonucleolytic activity of miRNasome-1. miRNA turnover assay performed in the absence or presence of 3 mM ATP (**top panel**) or AMP-PNP (**bottom panel**) with body-labeled *mir-84* and miRNasome-1, pre-incubated with antibodies as indicated. **(D)** Anti-XRN-2 antibody inhibits the endoribonucleolytic activity of miRNasome-1. Turnover assay performed in the absence of ATP with body-labeled *mir-84* and miRNasome-1, pre-incubated with antibodies as indicated. **(E)** ATP-independent formation of monoribonucleotides by recombinant XRN-2. 10 nM body-radiolabeled *mir-84* was subjected to *in vitro* miRNA turnover assay performed with purified recombinant XRN-2 (in an ascending concentration of 0.9 nM to 90 nM) or total lysate in the absence or presence of ATP, as indicated. The lower arrowhead indicates to monoribonucleotides (compare lanes 5, 8, 9). The reactions with the recombinant proteins were supplemented with 100 nM cold substrate miRNA (*mir-84*) to facilitate the monoribonucleotide formation. **(F)** Time-course experiment depicts gradual accumulation of the final 3-nt long products through progressive cleavage of the substrate miRNA at 3-nt intervals. Recombinant XRN-2 was incubated with body-radiolabeled *mir-84* in the absence of ATP. In this figure and the subsequent concerned figures, IgG purified from the rabbit pre-immune serum was used as control antibody. Representative images from three or more repeats have been presented.

Since the RNA employed above was body-radiolabeled and no significant amount of monoribonucleotide was generated, it was obviously not a product of an exoribonuclease activity, but endoribonuclease. Importantly, this 3-nt long product formation got heavily diminished, when we used the same substrate miRNA (5’-radiolabeled), where the phosphodiester bond between the 3^rd^ and 4^th^ nucleosides has been replaced with a PTO bond, and that essentially confirmed the endoribonucleolytic nature of the activity (**Fig. 3B bottom panel**). But, the same reaction yielded an additional band of 2-nt length, when the substrate miRNA was not an exact multiple of 3-nt, instead 23-nt long (*mir-237*, **Fig. 3B right panel**). These observations suggested that the endoribonuclease activity dices the substrate miRNA, beginning from the 5’-end, at 3-nt intervals in a highly processive manner. Therefore, by selecting the 24-nt long *mir-84* for these assays, we avoided simultaneous accumulation of multiple endoribonucleolytic products, and thus made the analysis simple and straightforward.

Interestingly, when the experiment described in **Fig. 3A** (**top panel**) was performed with the non-hydrolysable analog of ATP, AMP-PNP, it also exerted the same effect (**Fig. 3A bottom panel**), indicating that ribonucleotide binding and not its hydrolysis was required for the inhibition. Preincubation of miRNasome-1 with a polyclonal antibody against miRNasome-1.4, and not the other components, prevented both the ribonucleotides to exert their inhibitory effects (**Fig. 3C**), and thus suggested miRNasome-1.4 to be the ribonucleotide binding component, which mediates the inhibitory effect on the endoribonuclease activity of miRNasome-1 upon binding the ribonucleotide. Hence, it was apparent that although miRNasome-1.4 may have little influence on miRNA levels *in vivo* (**fig. S8**), but depending on the environment it might determine the modus operandi of miRNasome-1.

Although, none of the miRNasome-1 components were known and predicted to have any endoribonuclease activity (*2, 16, 24*), but preincubation of only anti-XRN-2 polyclonal antibody (*6, 27*) with miRNasome-1 could inhibit the formation of the 3-nt product, and thus suggested an endoribonuclease activity of miRNasome-1-residing XRN-2 on the substrate in the absence of ATP (**Fig. 3D**). We performed a super-shift assay and observed that the miRNP formed by miRNasome-1 and radiolabeled miRNA got further shifted up, upon incubation with the specific antibodies against the individual miRNasome-1 components (**fig. S11**). It excluded the possibility that antibody binding is leading to the dissociation of the miRNasome-1 complex, and accordingly the results obtained from the antibody-inhibition experiments are not fortuitous.

The intrinsic endoribonuclease activity of XRN-2 could be confirmed by checking the activity of the recombinant protein (**fig. S3A right panel**) on a body-radiolabeled *mir-84*, and it appeared to be highly processive as it diced the 24-nt long substrate into 3-nt long products (**Fig. 3E**). Notably, larger intermediate products could be detected only at earlier time points of the reaction, and their sizes indeed confirmed that the cleavage happens after every three nucleotides (**Fig. 3F**). We observed monoribonucleotide formation through exoribonuclease activity at higher protein concentrations (**Fig. 3E lane 5**), and lower concentrations only yielded endoribonucleolytic product (**Fig. 3E lanes 3, 4**), indicating a much stronger intrinsic endoribonuclease activity of XRN-2, over its exoribonuclease activity. ATP failed to exert any effect on either of the activities (**Fig. 3E lanes 6-8**). Monoribonucleotide formation was abolished, when the same assay was performed with a recombinant XRN-2 protein mutant for the exoribonuclease activity (D234-A, D236-A; reference *27*; **fig. S3A right panel**). However, the endoribonucleolytic product formation remained unchanged (**fig. S12 top arrowhead**), which confirmed that the active sites for these two activities are mutually exclusive. The recombinant versions of the other miRNasome-1 subunits didn’t show any nuclease activity (data not shown).

### miRNasome-1 is endowed with an ATP-dependent exoribonuclease activity and can switch between two alternative modes of action

The missing exoribonuclease activity of miRNasome-1 in the above reactions (**Figure 3A, C**), where it’s endoribonuclease activity was inhibited by ATP, could indeed be achieved upon supplementing the reaction with an increasing concentration of cold substrate miRNA. It led to the gradual increase in product accumulation (radiolabeled monoribonucleotides, **Figure 4A top panel**), which eventually got competed out by large excess of cold substrate in the reactions. This indicated to a possibility that in the concerned reactions described in the previous segment, the substrate concentration might have been way below the *Km* value of the exoribonuclease activity to yield any monoribonucleotide. Alternatively, it was also possible that the substrate concentration employed for the relevant reactions was significantly below the *K*_*d*_ value of the miRNA-binding receptor component of the miRNasome-1 complex, in case the exoribonuclease activity of the complex would be dependent on miRNA binding by such a dedicated receptor subunit. Interestingly, no monoribonucleotide was formed when ATP was replaced by AMP-PNP in this experiment (compare **Fig. 4A top and bottom panels**), suggesting that the conversion of the substrate into monoribonucleotides by miRNasome-1 is dependent on energy from ATP hydrolysis. Since, no monoribonucleotide was formed in the presence of AMP-PNP and optimum substrate concentration (**Fig. 4A bottom panel**), it could be suggested that energy might be required for the activation of the exoribonuclease active site of miRNasome-1-residing XRN-2.

**Fig 4.**
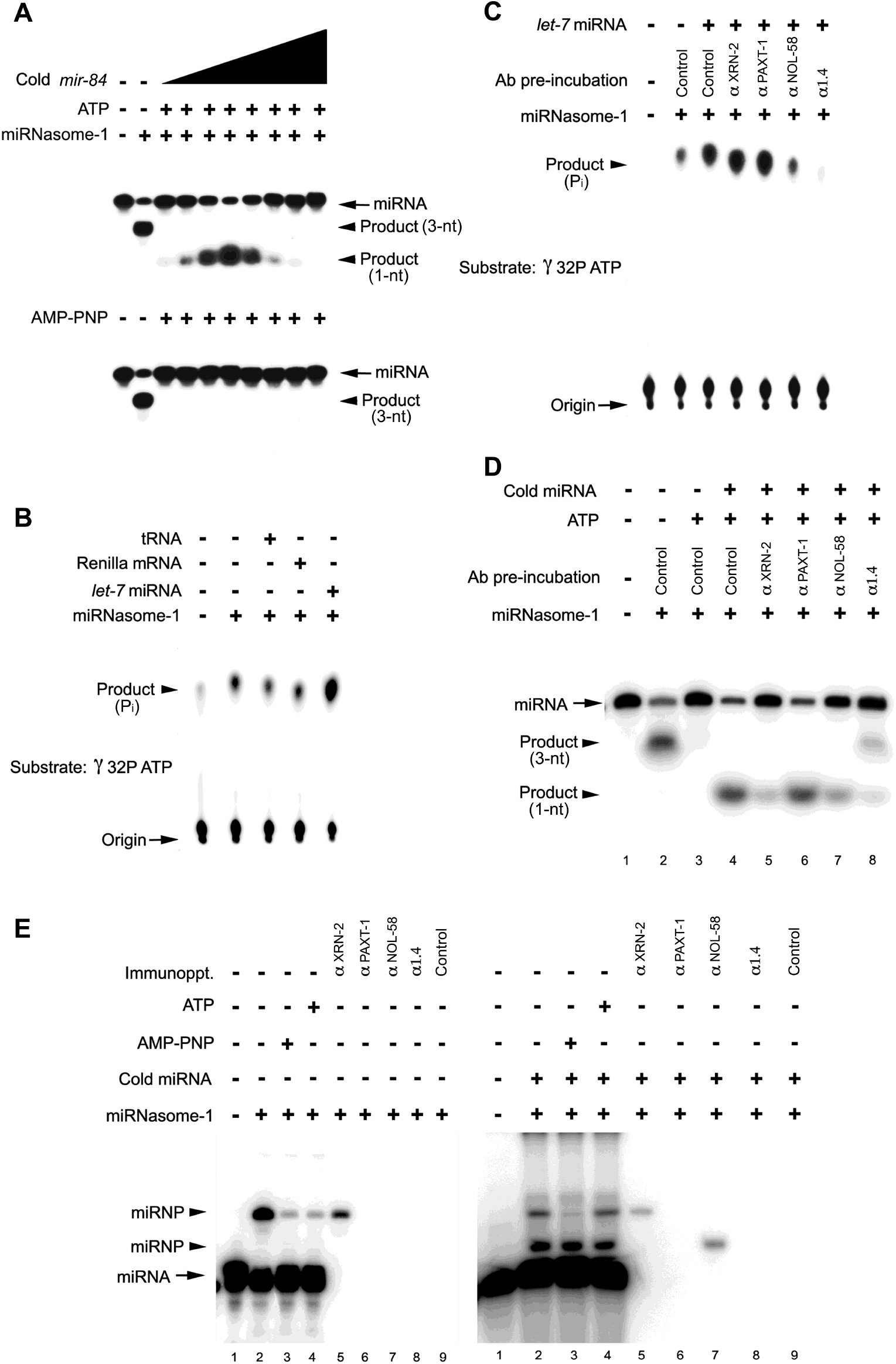
Energy from miRNA induced ATP hydrolysis is required for exoribonucleolysis of substrate miRNA by miRNasome-1. **(A) Top panel**. Purified miRNasome-1 (5 ng, ∼1.9 nM) was incubated with body-labeled mature *mir-84* (100 fmol, 10 nM) in the absence or presence of ATP (3.0 mM) as indicated. In order to achieve the exoribonucleolytic activity, unlabeled 5’-phosphorylated *mir-84* was added to the indicated reactions, in ascending concentration (25 nM to 1.6 µM). Lower arrowhead indicates to monoribonucleotides, which showed maximum accumulation in the presence of 100 nM cold substrate. **Bottom panel**. Reactions were performed as described above, except, ATP was replaced by the non-hydrolysable analog AMP-PNP. **(B)** ATPase activity of miRNasome-1 is specifically stimulated by miRNA. Purified miRNasome-1 was incubated with gamma-^32^P ATP in the absence or presence of indicated RNAs (100 nM). **(C)** Pre-incubation of miRNasome-1 with anti-NOL-58 antibody abolishes the stimulatory activity of miRNA, and anti-miRNasome-1.4 antibody completely abrogates the ATPase activity. Purified miRNasome-1 was incubated with gamma-^32^P ATP in the absence or presence of cold *let-7* miRNA (100 nM). **(D)** Anti-XRN-2 and anti-NOL-58 antibodies inhibit the formation of monoribonucleotides (bottom arrowhead). Turnover assay performed in the absence or presence of 3 mM ATP with body-radiolabeled *mir-84* (10 nM) and miRNasome-1 (∼1.9 nM), pre-incubated with antibodies as indicated. Cold synthetic *mir-84* miRNA (100 nM) was added to the indicated reactions to achieve the exoribonucleolytic activity of miRNasome-1. **(E)** Formation of miRNPs by miRNA [at low (10 nM, **left panel**) and high (110 nM, **right panel**) concentrations] and different components of miRNasome-1 in the absence or presence of ATP and AMP-PNP. In the absence (**left panel**) or presence (**right panel**) of 100 nM cold substrate-miRNA, IU-radiolabeled miRNA (10 nM) were incubated with miRNasome-1, pre-incubated without/ with ribonucleotides as indicated, followed by UV crosslinking, SDS-dissociation, and gel electrophoresis. The reactions in lane 2 (**for both the panels**) were performed in multiples, and subjected to immunoprecipitation using antibodies as indicated. Arrow indicates to free miRNA, and top arrowhead indicates to RNP comprising miRNA and XRN-2, and bottom arrowhead indicates to RNP comprising miRNA and NOL-58. Representative images from three or more repeats have been presented.

The observations made from **Fig. 4A** clearly indicated that miRNasome-1 might possess an intrinsic ATPase activity. Indeed, miRNasome-1 showed modest ATP-hydrolysis activity, which got significantly stimulated only in the presence of miRNA, and not tRNA or an mRNA (**Fig. 4B**). Notably, *let-7* showed a greater stimulatory effect than *lin-4*, and a vector backbone derived 30 nt long RNA, which indicated the miRNA mediated stimulatory activity to be a sequence specific event (data not shown). This stimulatory activity of the miRNA could be inhibited upon pre-incubation of the complex with anti-NOL-58 antibody (**Fig. 4C**). This clearly suggested that NOL-58 is probably the miRNA binding subunit, which stimulates the ATP hydrolysis. On the other hand, the antibody against miRNasome-1.4 abrogated miRNasome-1’s ATP hydrolysing activity (**Fig. 4C**). These results further suggested that miRNasome-1.4 of the complex harbors an ATPase activity, which gets stimulated through some conformational change relayed upon it, triggered through binding of miRNA by NOL-58. Accordingly, recombinant miRNasome-1.4 protein, and not the other subunits of miRNasome-1, showed an ATP hydrolysing activity (**fig S13A**), which, in expected lines, couldn’t be stimulated by miRNA, but did get inhibited with the antibody against miRNasome-1.4 (**fig. S13B)**. The ATP hydrolysing activity of miRNasome-1.4 could also be muted with orthovanadate, which is a structural mimic of phosphates, and thus, competes for phosphate binding sites in ATPases and phosphatases (**fig. S13C)**.

We have already observed that recombinant XRN-2 could produce monoribonucleotides in the absence of ATP (**Fig. 3E**). But, all the above results (**Fig. 4A-C**) indicated to a possibility that the miRNA getting converted into monoribonucleotides by the miRNasome-1 was not directly acted upon by the exoribonuclease active site of XRN-2, rather it was first bound elsewhere on the miRNasome-1, which could be NOL-58, and then it got transferred to the exoribonuclease active site of XRN-2 using the energy from the miRNA-stimulated ATP hydrolysis.

Consistent with the above hypothesis, in the presence of optimum concentration of substrate and ATP in a turnover assay, the monoribonucleotide formation by miRNasome-1 could be diminished not only by the anti-XRN-2 antibody, but also by the anti-NOL-58 antibody (**Fig. 4D, compare lanes 4, 5, 7**), thus indicating NOL-58 to be the possible receptor from where the RNA travels to the exoribonuclease active site of XRN-2. Interestingly, blocking of miRNasome-1 with the anti-miRNasome-1.4 antibody, and thereupon preventing ATP to bind to the ATP-binding subunit of the complex, not only reduced the formation of monoribonucleotides, but, at the same time it led to the vgeneration of the endoribonucleolytic product (**Fig. 4D, lane 8, top arrowhead**). This result was further supported, when we observed that in a miRNA turnover assay employing *miRNasome-1*.*4*(*RNAi*) lysate, despite the presence of ATP, there was a diminished monoribonucleotide formation and a concomitant accumulation of the endoribonucleolytic product, when the control lysate produced largely monoribonucleotides (**fig. S14**). Thus, it again demonstrated the mechanism of switching between two alternative modes of operation by miRNasome-1, depending on the availability of ATP.

### A putative travel-path of miRNA on miRNasome-1

To gain further mechanistic insights of the alternative modes of functioning of miRNasome-1, we needed to detect and identify the different miRNPs formed by miRNA (at low vs high concentrations) and the subunits of miRNasome-1 under different conditions (absence vs presence of ATP and AMP-PNP). We resorted to a binding experiment using miRNasome-1 and 5-Iodouridine-radiolabeled miRNA, followed by UV-crosslinking (*31, 32*), SDS dissociation of the non-crosslinked subunits and crosslinked-RNPs, and gel analysis. Immunoprecipitation reactions using antibodies against the individual miRNasome-1 subunits and crosslinked samples that were SDS-dissociated and then renatured (**Fig. 4E, left panel lanes 5-9 and right panel lanes 5-9**), revealed the identity of miRNasome-1 subunits (XRN-2, NOL-58; **Fig. 4E, left panel lane 5 and right panel lanes 5, 7**), involved in the formation of the major miRNP bands. We observed that reactions performed with low substrate concentration (10 nM) in the absence of any ribonucleotide, yielded a single major miRNP, comprising miRNA and XRN-2 (**Fig. 4E, left panel**). This essentially corroborated with the endoribonucleolytic activity of miRNasome-1 at low substrate concentration. More importantly, it indicated that XRN-2 of miRNasome-1 can directly catalyze endoribonucleolysis of the substrate, and doesn’t need any other subunit of the complex to mediate this action. In expected lines, this miRNP formation was diminished in the presence of ATP/ AMP-PNP (**Fig. 4E, left panel lanes 3, 4**), which corroborated with the inhibitory effect of ATP/ AMP-PNP on the endoribonuclease activity of miRNasome-1 (**Fig. 3A**).

We also noted that in the absence of any ribonucleotide, but at a high substrate concentration (∼110 nM, optimal for exoribonuclease activity), miRNA got crosslinked to XRN-2 and to a greater extent to NOL-58 (**Fig. 4E, right panel lane 2**), and that corroborated with endoribonucleolytic activity of XRN-2 on miRNA in the absence of ATP, and also reflected the binding of miRNA with the receptor component of the complex. However, addition of AMP-PNP diminished the interaction between the miRNA and XRN-2 (**Fig. 4E, right panel lane 3**), but showed the same robust crosslinking to NOL-58, which indicated that although the miRNA is bound to NOL-58, but not available for catalysis by any of the XRN-2 activities. Addition of ATP again brought back the miRNP formed by XRN-2, and showed slightly diminished formation of the RNP comprising miRNA and NOL-58, in comparison to that obtained in the presence of AMP-PNP. This modest diminishment (16% ± 2%) in RNP comprising miRNA and NOL-58 probably reflected the delivery of the NOL-58 bound miRNA to XRN-2, which might be due to a higher rate of miRNA transfer compared to the rate of miRNA-NOL58 RNP formation. Thus, it indicated that in order to have the NOL-58 bound miRNA transferred to XRN-2 to undergo exoribonucleolysis, energy from the hydrolysis of ATP is indispensable. However, we were unable to unravel the contribution of other subunits, if any, in this process of miRNA transfer, as a time-course of this crosslinking experiment was not fruitful, since, any shorter duration of UV-shining resulted in the formation of very week or no miRNPs (data not shown). Clearly, this macromolecular machine houses the repertoire to switch between two different (endoribonucleolytic vs exoribonucleolytic) modes of turnover mechanism through formation of distinct miRNPs, depending on the absence or presence of energy/ ATP (**Fig. 5**).

**Fig 5.**
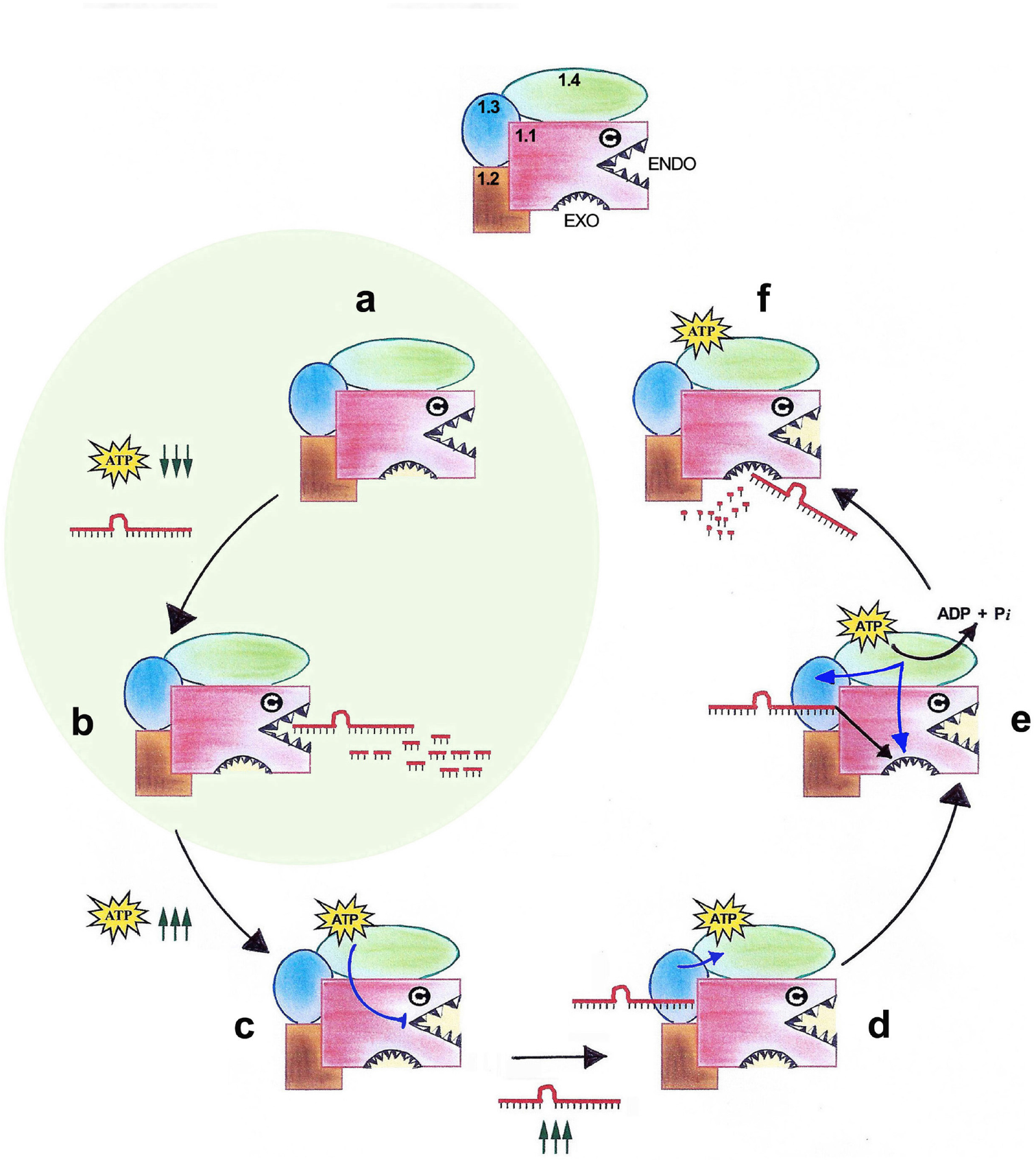
Model of miRNasome-1 *modi operandi* (anti-clockwise). The different subunits of miRNasome-1 (XRN-2 as 1.1, PAXT-1 as 1.2, NOL-58 as 1.3, miRNasome-1.4 as 1.4), and the endoribonuclease (ENDO) and exoribonuclease (EXO) active sites of XRN-2 are depicted on top. (**a, b**) The endoribonuclease activity acts on the miRNA more efficaciously than the exoribonuclease, the active site of which probably needs activation. (**b**) At low miRNA and ATP concentrations (depicted by downward arrows), turnover happens through the endoribonucleolytic mode. (**c**) ATP binding to miRNasome-1.4 exerts an inhibitory effect on the endoribonuclease activity of XRN-2. (**d, e**) At optimum concentration (depicted by upward arrows), miRNA binds to NOL-58, the RNA binding receptor, induces miRNasome-1.4, an ATP/ ribonucleotide binding protein, to hydrolyse the ATP, which further activates the exoribonuclease active site of XRN-2, and the NOL-58-bound-miRNA gets transferred to XRN-2 for exoribonucleolytic digestion. (**f**) Instantaneous reloading of ATP on miRNasome-1.4 would keep the endoribonuclease activity silent. ATP hydrolysis may also reinforce the inactive-state of the endoribonuclease active site. Blue lines indicate relaying of conformational changes. The mechanisms described in (**a)** and (**b)** (highlighted with a green circle) might be operating in dauer worms.

## DISCUSSION

Our results demonstrate that miRNasome-1 is indeed a multifaceted miRNA turnover complex, where two newly identified components (NOL-58 and miRNasome-1.4), govern the function of the ‘miRNase’-XRN-2. We also report a previously unknown endoribonuclease activity of the fundamentally important enzyme-XRN-2, and to our surprise, this activity is much more efficacious on the miRNAs than the previously known exoribonuclease activity. ATP binding to miRNasome-1.4 exerts an inhibitory effect on this endoribonuclease activity. Whereas, miRNA binding to NOL-58, the *bona fide* RNA-binding subunit, stimulates ATP hydrolysis, and the energy is utilized towards the transfer of the miRNA to the ‘miRNase’ for its exoribonucleolysis (**Fig. 5**).

Perturbation of NOL-58 leads to the specific accumulation of a number of mature miRNAs involved in several important cellular and developmental processes like cell proliferation and differentiation (*let-7*), muscle development (*mir-1*), neuronal metabolism (*mir-124*) etc. A greater magnitude of accumulation of mature miRNAs in NOL-58 depleted samples, compared to XRN-2 depleted samples, was observed (**Fig. 2A, fig. S8**), which was probably due to a more efficient knockdown of the former (**fig. S9**), under the employed conditions. Notably, drastically different affinities of the recombinant NOL-58 for good-substrate and poor-substrate miRNAs (eg. *let-7* vs *lin-4*, data not shown), suggested that the intrinsic affinity of NOL-58 towards different miRNAs might be playing a key role in conferring the specificity to miRNasome-1. However, perturbation of miRNasome-1.4 didnt lead to any accumulation of the cohort of miRNAs, we checked. It is most likely that if miRNasome-1.4 would be absent or inhibited to bind ATP, the miRNasome-1-residing XRN-2 would still be able to act endoribonucleolytically on the substrate miRNAs (**Fig 3C, Fig 4D**), which would be sufficient to inactivate those miRNAs. Accordingly, **fig. S14** clearly demonstrated that miRNasome-1.4 depleted lysate results in the accumulation of the endoribonucleolytic product by cleavage of the intact miRNA in a miRNA turnover assay, even when it is performed in the presence of ATP. Thus, although, miRNasome-1.4 is required for the functionality of the miRNasome-1 complex, but its perturbation may not lead to aberrant miRNA accumulation. Notably, monoribonucleotides produced from energy-dependent exoribonucleolysis of miRNAs in continuously growing worms are likely to recycle back readily to the biosynthetic pathways, and thus may be preferred over 3-nt long products. However, physiological significance, if any, of the 3-nt long product of the highly processive energy-independent endoribonuclease activity of miRNasome-1 is yet to be unraveled. Nevertheless, it is not very difficult to imagine the 3-nt product as a sub-cellular signal allowing the organism to sense its environment and re-calibrate its metabolism accordingly.

Exoribonuclease activity of XRN-2 was characterized using endogenous proteins purified from human samples and yeast (*33, 34*), and very little or no endoribonuclease activity was reported (*34*). It is possible that the selection of the very short (4-nt) and very long substrates capable of forming complex structures, and the heterogeneity of the employed protein excluded the detection of the endoribonuclease activity of XRN-2 (*34*), especially now, when we know of an example, where a protein (miRNasome-1.4) inhibits the endoribonuclease activity of XRN-2, upon binding to ribonucleotides. Our studies reveal that the endoribonuclease activity of XRN-2 acts efficiently on unstructured RNAs, and requires a free 5’-unpaired-end, and accordingly, structured tRNA with a 5’-basepaired end proved to be a very poor substrate (**fig. S4B**). Therefore, selection of unstructured short miRNAs as substrates indeed helped us to detect this previously unknown activity. Furthermore, recombinant XRN-2 protein also shows a very weak exoribonuclease activity on miRNAs, compared to its endoribonuclease activity. A closer look at the available worm XRN-2 structure reveals that the entry-point for XRN-2’s exoribonuclease active site tunnel is partially occluded by a bulky residue (W670, reference *35*). Thus, it is possible that an *in vitro* reaction performed with high substrate and protein concentrations led to effective collisions, which landed the substrate miRNA in the exoribonuclease active site tunnel (**Fig. 3E**). It is also possible that the ATP-dependent exoribonuclease activity of miRNasome-1 on miRNA involves conformational changes to allow the substrate reach XRN-2’s exoribonuclease active site more readily.

Overall, miRNasome-1 appears to be a potent molecular machine, involved in the maintenance of physiological levels of a number of important miRNAs, and hence it crucially contributes to target mRNA homeostasis and normal worm development. Current literature indicates that under normal circumstances the subunits of this complex are nuclear localized (*2, 6, 24, 36, 37*). This is in contrarian to the expectation that miRNasome-1 should be in the same cytoplasmic compartment as miRISC to modulate its activity. It has been suggested before that a miRNA residing in miRISC must undergo a ‘release step’ before its turnover can happen (*5*). Therefore, a miRNA may get dislodged from the miRNA-Argonautes in the cytosol by the ‘release factor’, before its transportation to the nucleus for degradation. Moreover, since, it has already been shown that miRNA-Argonautes can be imported into the nucleus (*38, 39*), it is also possible that a mechanism operates to actively remove the ‘destined miRNA-Argonautes’ from the cytosolic workbenches, so that their miRNAs can undergo degradation.

Interestingly, *in vitro* assays revealed that the XRN-2 dependent exoribonuclease activity on miRNAs appears to be much stronger in the total lysate (**Figure 1A**), in spite of a lower abundance of XRN-2, compared to the activity in the isolated system as observed in **Figures 1F** and **4A**, where XRN-2 is much more abundant in the purified miRNasome-1. Although, we can not completely exclude the possibility that our purified complex is devoid of a component, which facilitates substrate binding or increases its affinity for the substrate, but it is also possible that *in vivo* miRNasome-1 does not act directly on miRNAs, rather the substrate is delivered to it by some other carrier protein, working upstream in the miRNA turnover pathway. That can possibly be the candidate, which ‘releases’ the miRNAs from the grasp of miRNA-Argonautes. Therefore, such miRNA bound carrier protein (miRNP)-miRNasome-1 interaction might play a crucial role in modulating the affinity of the receptor protein (NOL-58) in the receiving complex (miRNasome-1). At any rate, miRNasome-1 emerges to be an important ‘unit’ of the miRNA turnover machinery. Notably, interactions between such ‘units’ could well achieve the stepwise mechanism of the miRNA turnover pathway observed in the worms (*5*).

Our *ex vivo* studies with miRNasome-1 demonstrated that this biological machine is highly receptive of its environment, and depending on the availability of energy it can operate in two alternative modes of catalytic mechanism, where its energy-independent endoribonuclease activity is much more efficacious than that of the energy-dependent exoribonuclease activity. Understandably, such an ATP-independent endoribonucleolytic-mode of turnover mechanism would be highly suitable to withstand a challenging environment, where food is scarce and thus energy needs to be conserved. One such physiologically relevant situation might be the miRNA homeostasis during the quiescent dauer stage (*40*), where cellular ATP and other high-energy phosphates are known to get diminished than that of the normal stages of life cycle (*41*), and thus potentially may have played a crucial role in the adaptation and survival of worms. It has also been noticed that the levels of some of the potential miRNasome-1 substrate miRNAs (eg. *mir-241, mir-237* etc) in the dauer stage are significantly reduced compared to that observed in the relevant stages of the continuously growing worms (L2/ L3, references *42, 43*). Accordingly, that justifies the higher efficacy of the endoribonuclease activity of miRNasome-1 over its exoribonuclease activity, which might get employed during the dauer stage (**Fig. 5**).

Finally, we have identified two previously unknown enzymatic activities, ATPase activity of miRNasome-1.4 and endoribonuclease activity of XRN-2. miRNasome-1.4 (gene number B0024.11) has been described as a homolog of human PUS7, and it does not possess a P-loop or Walker-A/B motifs characteristic of the standard catalytic cores of ATPases. Therefore, characterization of its active site and the mechanism of catalysis, and further determination of its *in vivo* roles would certainly accentuate our understanding. Similarly, identification of the XRN-2 endoribonuclease active site, its mechanism of action, and deciphering its *in vivo* roles, especially in quiescent dauer worms might reveal some interesting biology. This is particularly significant considering the close analogy of dauer quiescence to that of the quiescent adult stem cells in higher eukaryotes and mammals (*43, 44*), and since, the dauer worms are analogous to the infective stage of the parasitic nematodes (*40*). Therefore, understanding of miRNasome-1 function under different physiological conditions would not only help us to understand dynamic miRNA metabolism, but might also create a platform for future translational research.

## ACKNOWLEDGMENTS

We thank Helge Grosshans and Craig C. Mello for worm strains and reagents. We are indebted to Chris D. Lima for advises on *in vitro* reconstitution of macromolecular complex, Moumita Sardar, Srinivas Babu, and late Monika Fasler for technical help. This work has been supported by the Wellcome Trust (UK) - DBT (Govt. of India) India Alliance through a fellowship to SC. MS, obtained fellowship from the University Grants Commission (Govt. of India). ARW, PK received support from SERB/ DST (Govt. of India).

## AUTHOR CONTRIBUTIONS

SC designed the research and experiments. MS, ARW, PK performed the experiments. MS, ARW, PK and SC together analyzed the results, and SC wrote the manuscript.

