## Supplementary Information for "A multifaceted microRNA turnover complex from *Caenorhabditis elegans*"

**This file includes:**
Materials and Methods
Supplementary Text
Figs. S1 to S14 with their captions
Table S1

#### Materials and Methods

**Worm strains, *let-7(n2853)* suppression and RNAi.** The wild type strain *C. elegans* var. Bristol strain *N2* has been used. The other four strains were *let-7*(*n2853*) (a temperature sensitive strain hosting a single G-A point mutation in the seed sequence of *let-7* miRNA; reference *25*), *gfp::alg-1; gfp::alg-2* (strain expressing integrated multicopy array of *gfp*-tagged versions of both *C. elegans* miRNA argonautes, *alg-1* and *alg-2*, under the control of endogenous promoters and 3'-UTRs; reference *30*), *xrn-2ts* [*xrn-2* null mutant, *xrn-2*(*tm3473*), recovered with an integrated single copy *xrn-2* transgene with P107-L mutation conferring temperature sensitivity; reference *27*]. Suppressors of *let-7*(*n2853*) were identified by RNAi through feeding, where worms were grown on RNAi plates at 25°C, essentially as described before (*29*), and any minor modifications have been mentioned in appropriate places. Suppression assay results were an average of live worms (where n = 200), 50 hrs post plating from L1 developmental stage, from three biological replicates ± SEM. All other phenotypic readouts were also an average from three biological replicates ± SEM, where n = 200 (except for studies on alae structure, where n = 50). Except for *paxt-1*, full-length cDNAs of the respective genes were cloned in *L4440* vector to prepare the RNAi clones. RNAi clone for *paxt-1* was used from GE Healthcare/ Dharmacon *C. elegans* RNAi Collection.

**RNA isolation, Northern blotting, RT-qPCR.** Total RNA was isolated from staged L4 worms, using Trizol (Ambion) method as described before (*45*). Northern blotting of endogenous RNA was done as described (*46*). 5'-labeled DNA oligos were used as probes, where OptiKinase™ (Phosphatase minus mutant of T4 polynucleotide kinase [PNK, Affymetrix]) and γ-$^{32}$P-ATP were used to perform the 5'-end labeling. The hybridizations for the members of the *let-7* family of miRNAs were carried out at an elevated temperature of 40°C in order to minimize the binding of the probe to unintended *let-7* sisters (*5*). For northern analysis of *in vitro* pre-*let-7* processing assay products (*5*), the lysates were pre-treated with micrococcal nuclease (MN, NEB, 2.0 µl (4000 Gels Units)/100 µg of lyaste) for 10 min at 37°C, followed by addition of EGTA to a final concentration of 7.5 mM. MN pre-treatment was done to digest all endogenous RNAs from the lysate, and thus ruling out any possibility of detection of endogenous RNA. Excess EGTA was used to terminate the MN treatment through chelation of $Ca^{++}$. '– EGTA' lysate served as a positive control for MN activity that cleared all the RNA, including exogenous RNA, resulting in complete loss of signal. Following incubation of RNAs in the lysates, the samples were phenol-chloroform extracted and alcohol precipitated in the presence of glycogen (20 µg/ reaction, Roche). The recovered samples were subjected to northern probing as per conditions stated above. For RT-qPCR, 2 µg of total RNA of the respective samples, was reverse transcribed using SuperScript™ III Reverse Transcriptase (Invitrogen) in 20 µl reaction containing 5 mM DTT, 0.5 mM dNTPs, 2.5 µM oligo $(dT)_{20}$ and 1 µl of enzyme, in accord with the manufacturer's protocol. 1.5 µl of each of the reverse-transcription reactions were amplified with PowerUp™ SYBR Green Master Mix (Applied Biosystems) in a total volume of 10 µl using specific forward and reverse primers at a concentration of 0.5 µM each, and analyzed in a ViiA™ 7 Real-Time PCR System using ΔΔCt method. The primers used are described below. Values were normalized against the value of act-1 expression. The experimental values, representing means (±SEM) from three independent biological replicates, were compared to the expression levels of the corresponding *let-7*(*n2853*)/ *N2* (Empty Vector/ Control) samples, which were set as 1.

**Cloning and expression of recombinant XRN-2, PAXT-1, NOL-58, miRNasome-1.4.** cDNA was generated from *C. elegans* total RNA using SuperScript™ III Reverse Transcriptase (Invitrogen) and oligo $(dT)_{20}$. Then *xrn-2* cDNA was PCR amplified from the cDNA using gene specific primers and Q5 High-Fidelity DNA Polymerase (NEB), and cloned in a TOPO TA vector (Invitrogen). The sequence confirmed correct ORF was subcloned in pFastBac$^{TM}$ HT B (Invitrogen), and further used to generate the recombinant bacmid DNA as per the manufacturer's protocol (Bac-to-Bac Baculovirus Expression System, Invitrogen), towards expression of the recombinant N-terminal 6XHis tag containing protein in insect cells (Sf9). Sf9 cells infected with P2 viral stock at a MOI of 5 were harvested 52 hrs post-infection. The cells were lysed using a hypotonic lysis buffer and the intact nuclei were subjected to high salt treatment to rupture the nuclear membrane. The nuclear supernatant was subjected to PEG 8000 treatment to precipitate the DNA, and the cleared supernatant was used to charge an affinity column with Ni-NTA resin (Qiagen), and the recombinant protein was eluted as per the manufacturer's protocol and as described (*47*). The same expression system was also used to express N-terminal Strep tag containing wild type and exoribonuclease mutant (D234-A, D236-A) XRN-2 proteins, which were purified using Strep-Tactin$^{\mathbf{R}}$XT Superflow$^{\mathbf{R}}$ high capacity resin (IBA, Germany). Of note, high salt (0.25-0.5M) containing wash and elution buffers were used for the removal of nonspecific host proteins and for the complete elution of the recombinant protein, respectively.

*paxt-1*, *nol-58*, *miRNasome-1.4* cDNAs were also generated as described above and cloned in pGEX-4T-1 vector (GE Healthcare). N-terminal GST-tagged proteins were purified from BL-21(DE3)/ Rosetta™ cells using Glutathione Sepharose™ 4B resin (GE Healthcare) as per the manufacturer's protocol. Notably, high salt (0.25-0.75M) containing wash and elution buffers were used for the removal of nonspecific host proteins and for the complete elution of the recombinant proteins, respectively. All the above proteins were subjected to dialysis to remove the excess salt before performing any functional assay.

**Generation of recombinant mutant XRN-2 protein with substituted amino acids.** Point mutations (D234-A, D236-A) were introduced in the full length *C. elegans xrn-2* gene through site directed mutagenesis, following the protocol detailed in the QuikChange site directed mutagenesis kit (Agilent), and sequence confirmed.

**Preparation of RNA substrates.** Mature *let-7,* mature *mir-84*, and mature *mir-237* RNA were prepared as per the methods described before (*5, 48*). In brief, a chimeric RNA comprising of a hammerhead ribozyme in its 5'-end followed by the mature *let-7/ mir-84/ mir-237* sequence was transcribed from DNA cassettes using a MEGAshortscript™ T7 transcription kit (Ambion) in presence of α-$^{32}$P-UTP (and 5-Iodouridine), as per the supplier's protocol. The DNA cassettes were prepared by annealing of appropriate forward and reverse primers (please see the oligo sequence section) followed by fill-in reactions using Klenow (NEB). Double stranded DNAs of appropriate length were gel purified and directly used as template for *in vitro* transcription. The gel purified products were also PCR amplified, cloned and sequence confirmed. While transcription reaction is ongoing, self-processing of the ribozyme-containing transcripts occur simultaneously.

The resulting mature *let-7/ mir-84 /mir-237* containing 5' hydroxyl groups were size purified by 7 M urea/ 8-10% PAGE. After recovery RNAs were 5' phosphorylated by T4 PNK (Affymetrix/ NEB) and ATP.

5' radiolabeling of synthetic pre-*let-7*, mature miRNAs (Microsynth AG, Switzerland) and tRNA (after dephosphorylation, yeast $tRNA^{Phe}$, Sigma) were done using OptiKinase™ T4 PNK (Affymetrix) and $\gamma$-$^{32}$P ATP. Before use the radiolabeled synthetic pre-*let-7* RNA was subjected to refolding as described before (*48*). 3' radiolabeling and blocking of synthetic mature miRNAs were done using T4 RNA ligase (Ambion) and [5'-$^{32}$P] pCp, according to manufacturer's instructions. All RNAs were size purified using 7 M urea/ 8-10% PAGE.

The 5'-7-methyl-G-capped RL reporter mRNAs (*Renilla* luciferase with an artificial 3'-UTR harbouring 3X Bulged *let-7* complementary sites; reference *5*), were prepared through *in vitro* run-off transcription of appropriately digested plasmids using reagents from a MEGAscript T7 Transcription kit and cap analog $m^7G(5')ppp(5')G$ from Ambion, as per the manufacturer's instruction. After phenol-chloroform extraction and alcohol precipitation the RNAs were polyadenylated using *E. coli* Poly(A) polymerase (NEB) and ATP. The above template was also used to synthesize a long transcript/ ORF through *in vitro* run-off transcription, using a MEGAscript T7 Transcription kit.

**Preparation of worm lysate.** Staged L4 worms were grown on plates and harvested with M9, and washed thrice with the same buffer to eliminate any bacteria. The worm pellet was then resuspended in extraction buffer (25 mM HEPES [pH 7.4], 2.5 mM DTT, 2.5 mM $MgCl_2$, 0.1% Triton-X 100, 100 mM KCl, 10% Glycerol, 1X SIGMAFAST™

Protease Inhibitor Cocktail) and ground in liquid $N_2$. The clear supernatant was collected after spinning the thawed sample at 16,000 g for 30 min, which was further sequentially passed through Miracloth (Calbiochem), empty Poly-Prep Chromatography Column (Bio-Rad), and 0.4 micron filter unit (Millipore), and designated as 'total lysate/ worm lysate' or simply 'lysate'. Of note, cleared worm lysate was used in all the *in vitro* assays involving lysate.

**Co-immunoprecipitation.** Cleared worm lysate was prepared as described above, and co-immunoprecipitations on samples with/ without RNaseA treatment were essentially performed following the methods described before (*6*). Of note, 3 mg sample and 5 ug of purified antibody was used for each immunoprecipitation reaction, and antibodies were crosslinked to protein G magnetic beads (PureProteome$^{TM}$ Protein G Magnetic Beads, Millipore).

***in vitro* turnover assay/ pre-*miRNA* processing assay.** Radiolabeled RNAs (pre-*let-7*/ mature *let-7*/ mature *mir-84,* tRNA, ORF [transcript of an Open Reading Frame], mRNA) approximately 100 fmol, if not otherwise indicated, were incubated with cleared worm lysate or a given chromatographic (Capto Q) fraction (1-5 μg) in 1X **A**ssay **B**uffer (AB; 25 mM HEPES [pH 7.4], 2.5 mM DTT, 5 mM $MgCl_2$, 100 mM KCl, 2.5 mM ATP) in a volume of 10 μl at 25°C for 15 min. Turnover assays with mature miRNA, tRNA, mRNA, and 100 ng of purified miRNasome-1 were performed in the AB same as above, except with a higher ATP concentration (7.5 mM). Turnover assays with mature miRNA, tRNA, mRNA and recombinant XRN-2 (1-100 ng) were also performed in the AB containing 7.5 mM ATP or as indicated. The reactions were terminated by addition of 1X volume formamide gel loading buffer (95% Formamide, 0.2% SDS, 1 mM EDTA, 0.04%

Xylene Cyanol, 0.04% Bromophenol Blue) followed by heating at 65°C for 5 min and chilling on ice. Equal volumes of the samples were then subjected to 7 M urea/8-16% PAGE followed by gel drying and autoradiography or phosphor-imaging.

The RNA size ladders for *let-7, mir-84* and *mir-237* were generated as per the Ambion's partial alkaline hydrolysis protocol, where a 5'-radiolabeled RNA (*let-7, mir-84, mir-237*) was subjected to incomplete hydrolysis by alkali.

**miRNA-protein (miRNasome-1 subunits) co-immunoprecipitation.** 1 pmol of 5'-32P-radiolabeled and 3'-biotinylated *let-7* miRNA, where all the phosphodiester bonds have been replaced by phosphorothioate (PTO) bonds to increase the stability of the RNA, was incubated with 100 µg of total worm lysate in a volume of 150 µl on ice for 15 minutes. The aforementioned reaction was performed in multiples, and after incubation the reaction volumes were increased to 450 µl with 1X AB and subjected to immunoprecipitation at 4°C for 2-3 hrs using antibodies against miRNasome-1 subunits, IgG purified from pre-immune serum (Control), and Protein G Sepharose™ 4 FastFlow (GE healthcare). The recovered sepharose beads were subjected to phenol-chloroform extraction and alcohol precipitation in the presence of glycogen (20 µg/ reaction, Roche). Finally, the recovered RNA samples were subjected to urea PAGE analysis.

**Coupled pre-*let-7* processing and Argonaute immunoprecipitation.** Pre-*let-7* processing assay was performed as described above using lysate obtained from a strain in which both the *C. elegans* miRISC Argonaute proteins ALG-1 & ALG-2 are tagged with GFP ('GFP/AGO'; reference *30*). After incubation for 15 min at 25°C, the reaction volumes were increased to 200 µl with 1X AB and subjected to immunoprecipitation at

4°C for 2-3 hrs using an α-GFP antibody (α-GFP mouse IgG; monoclonal antibody, Roche [Cat. # 11 814 460 001]) and Protein G Sepharose™ 4 FastFlow (GE healthcare). The recovered sepharose beads were suspended in formamide gel loading buffer (95% Formamide, 0.2% SDS, 1 mM EDTA, 0.04% Xylene Cyanol, 0.04% Bromophenol Blue), heated at 65°C for 5 min, spun briefly and the supernatants were subjected to urea PAGE analysis (*5*). The post-immunoprecipitate supernatants were also recovered through phenol-chloroform extraction and alcohol precipitation in the presence of glycogen (20 μg/ reaction, Roche), and subjected to urea PAGE analysis.

**Fractionation of staged and cleared worm lysate.** 500 mg to 1.0 gm of staged and cleared L4 worm lysate was adjusted to a protein concentration of 3-5 mg/ ml, and $MgCl_2$ concentration of 5.0 mM, and charged on a custom made Tricorn™ High Performance column (GE Healthcare) packed with ~ 8.0 ml of Capto Q resin (GE Healthcare), using an AKTA™ avant FPLC system. After sample loading, the column was extensively washed with 1X assay buffer (excluding ATP), and subjected to elution in the same buffer, but with a continuous ascending salt gradient (150 mM to 500 mM). The different fractions were subjected to SDS PAGE, followed by western blotting using anti-XRN-2 antibody. Only XRN-2 containing fractions were subjected to RNA-affinity purification. Notably, the XRN-2 containing fractions from the Capto Q column could also be further fractionated in a Resource Q column, and the XRN-2 containing fractions were subjected to RNA-affinity purification. This additional step modestly increased the final yield, without affecting the properties of the purified complex, and thus, considered non-essential.

**RNA-affinity purification.** A 5'-phosphorylated *let-7* having all its phosphodiester bonds replaced with phosphorothioate bonds, and the 3'-end of the miRNA connected to a biotin through a $C_{18}$ alkyl group was used as the affinity arm (Microsynth AG, Switzerland) for RNA-affinity purification. 100-200 nmol of the aforementioned affinity arm was immobilized on Streptavidin Sepharose™ High Performance beads (GE Healthcare), and incubated with the relevant Capto Q fractions in 1X assay buffer having the final salt concentration adjusted to 500 mM, at 4°C for 30 minutes on a rotator. This high salt incubation essentially excluded binding of non-specific RNA-binding proteins and other miRNA-interacting proteins, including miRNA-Argonautes, to the affinity arm. The protein-bound beads were settled in a Poly-Prep Chromatography Column (Bio-Rad), washed with 1X assay buffer with increasing concentrations of salt (500 mM to 750 mM KCl), and finally subjected to elution with 1X assay buffer containing high salt (1.0 M KCl). The eluate was dialyzed to bring down the salt concentration to an approximate final concentration of 100-150 mM, and subjected to 5% native or native-gradient (4-16% or 4-20%) gel electrophoresis followed by Coomassie Brilliant Blue/ silver staining (SilverQuest™ Staining Kit, Invitrogen) or western blotting. Notably, at times, we did obtain a few minor additional bands, and in those cases gel filtration chromatography was employed to obtain the pure high molecular weight complex. Western blotting was performed employing one-fifth to one-tenth of the protein used for gel staining. For determination of molar ratio of the constituent subunits of miRNasome-1, the purified protein complex was resolved on a SDS-PAGE and stained with Coomassie Brilliant Blue. The densities of the individual subunits were measured using GS-900™ software (Bio-Rad), followed by normalization with their respective molecular weights. The normalized densities of the different subunits showed a 1:1:1:1 ratio with respect to each other.

**Antibody inhibition assays.** miRNasome-1 (5 ng) was incubated with 25-50 ng of IgG, purified from the concerned rat/ rabbit polyclonal antiserum in 1X assay buffer without ATP on ice for 30 minutes with occasional gentle mixing, followed by addition of radiolabeled miRNA and other relevant reagents to a final volume of 10 μl. The reactions were incubated at 25°C for 15 minutes and then terminated as described above, and subjected to 7 M urea/10-15% PAGE followed by gel drying and autoradiography or phosphor-imaging. It is to be noted that in the concerned figures (Figures 1, 3, 4), α-miRNasome-1.4 antibody has been represented as α-1.4. IgG purified from the rabbit pre-immune serum has been represented as Control.

**ATPase assay.** 5 ng of miRNasome-1 or 10-20 ng of recombinant miRNasome-1 subunit proteins were incubated with 2-4 μCi of $\gamma$-$^{32}$P-ATP (Specific Activity: 4000 Ci/ mMol) in the presence of 3.0 mM of cold ATP at 25ºC for 15 minutes. Different cold RNAs (miRNA, tRNA, mRNA), each at a final concentration of 100 nM, were added to individual reactions to check their effector activities.

**Thin Layer Chromatography (TLC).** Mature miRNA turnover reactions, using 5'-PNK radiolabeled or $\alpha$-$^{32}$P-UTP labeled miRNA, were stopped by the addition of SDS to 1% and EDTA to 10 mM. 1μl aliquots of each reaction were spotted onto pre-washed PEI cellulose plates (Sigma/ Millipore) and developed sequentially with 0.5 M LiCl and 1.0 M formic acid (*5*). ATPase assay reactions were also terminated as above, and similarly spotted on pre-washed PEI cellulose plates (Sigma/ Millipore) and developed sequentially with 0.1 M acetic acid and 1.0 M LiCl as described before (*49*), and subjected to autoradiography or phosphor-imaging.

**Crosslinking and immunoprecipitation assay.** miRNasome-1 (5 ng) were incubated with 5-IU-radiolabeled *mir-84* (10 nM), in the presence of cold *mir-84* (100 nM) for 2-5 min on ice, after a pre-incubation with the AB containing the indicated ribonucleotides (3 mM) for 2-5 min, and irradiated with 365-nm UV light in a UVP cross-linker (Cambridge, UK) for 10 min with cooling to form RNA-protein crosslinks (*31, 32*). Then SDS was added to a final concentration of 0.2% to dissociate the different RNPs from the proteins of the complex which didn't participate to form RNPs. These reactions were either directly used for gel analyses or diluted 20-fold into a renaturation buffer (1X TETN 250: 250 mM Tris-HCl [pH 7.5], 5 mM EDTA, 0.5% Triton X-100, 250 mM NaCl, 2 mg of BSA/ml) and incubated on ice for 45 min, followed by addition of 100 ng of IgG purified from the concerned rat/ rabbit polyclonal antiserum or rabbit pre-immune serum. Immunoprecipitation was performed with Protein G Sepharose™ 4 Fast Flow (GE Healthcare). Immune complexes were dissociated in formamide sample buffer containing 0.5% SDS, and resolved through 7 M urea–9% PAGE (Acrylamide: Bis-acrylamide = 39: 1) in the presence of 0.05% SDS.

**Antibodies and Western blotting.** Bacterially expressed pure full length NOL-58 and miRNasome-1.4 were injected in rabbit (New Zealand White Rabbits) as per the methods described before (*50*). The collected sera were subjected to affinity purification using Protein A Sepharose™ CL 4B (GE Healthcare). IgG was eluted using Pierce™ Gentle Ag/Ab Elution Buffer, pH 6.6, dialyzed against 1X assay buffer (without ATP), and concentrated using Amicon Ultra (Millipore). Rat α-XRN-2 serum, rabbit α-XRN-1 polyclonal IgG, and rabbit α-PAXT-1 polyclonal IgG were generously provided by Helge Grosshans (*27, 8, 6*).

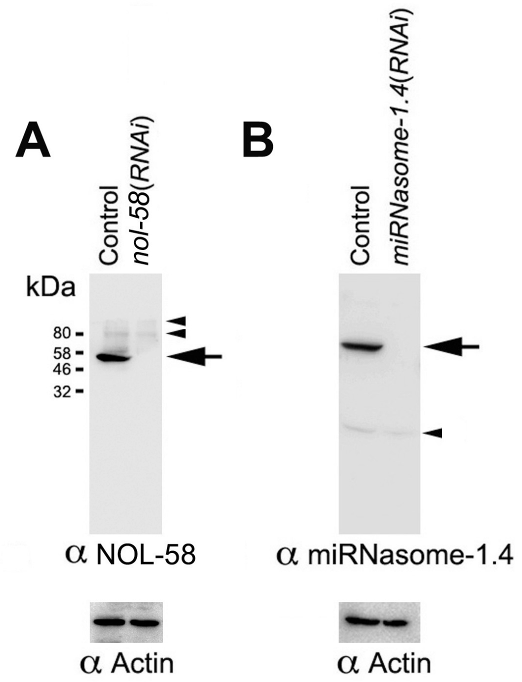


**Specificity of NOL-58 and miRNasome-1.4 antisera**

Western blotting identified single bands (arrow) of desired size from total worm lysate using α-NOL-58 (**A**) and α-miRNasome-1.4 (**B**) sera, which got heavily diminished upon RNAi depletion of the respective gene products, as indicated. Minor non-specific bands are indicated with arrowheads. Actin served as loading control.

For western blotting, α-NOL-58 and α-miRNasome-1.4 sera were used at 1: 2000 dilutions, followed by the use of α-Rabbit HRP-conjugated secondary antibody (GE Healthcare) at a dilution of 1: 10,000. Detection was performed with Amersham™

ECL™ Prime Western Blotting Detection Reagent, and Amersham Hyperfilm™ ECL or ImageQuant LAS 4000 Chemiluminescence Imager (GE Healthcare). Band sizes (relative molecular mass) were determined using GS-900™ software (Bio-Rad).

**Electrophoretic mobility shift assay.** miRNA binding activity of the endogenous miRNasome-1 was assayed by incubating the purified complex with 5'-PNK labeled and PTO stabilized *let-7* in AB for 30 min on ice, followed by observing and confirming the electrophoretic mobility shift of the RNP complex in a 6% polyacrylamide gel (59:1), resolved at 4°C in the presence of 5 mM $MgCl_2$. To perform super-shift, the relevant antibody was added to a given reaction after 30 min of only RNA and protein incubation, and further 30 min incubation was carried out before gel analysis. The first-four phosphodiester bonds from the 5'-end of the substrate (*let-7*) miRNA were replaced with PTO bonds.

**Reconstitution and purification of miRNasome-1.** Purified recombinant GST-tagged PAXT-1, NOL-58, and miRNasome-1.4 expressed in *E. coli*, and 6XHis tagged XRN-2 expressed in insect cells were mixed in equimolar ratio in AB containing no ATP and high salt (350 mM KCl), and subjected to two-step dialysis as described (*51*). *in vitro* reconstitutions were carried out in a volume of 1.0 ml using ~2.0 mg of total protein. The post-dialysed samples were subjected to affinity purification using Glutathione Sepharose™ 4B resin, followed by Gel Filtration Chromatography (GFC, Superdex 200 Increase 10/300 GL [GE Healthcare]) to separate the *bona fide* complex from sub-complexes and individual proteins.

**Mass-spectrometry.** Sample was subjected to in-gel digestion using Trypsin Gold (Promega). Digested peptides were reconstituted in 15 µl of 2% CAN with 0.1% formic acid. 1 µl of the sample was injected in a chromatography column, where it was subjected to a RPLC gradient for 70 minutes, followed by acquisition of the data on a connected LTQ-Orbitrap Discovery MS (Thermo Scientific). Identities of the peptides were searched using MASCOT 2.4 on Swiss-prot, TrEMBL and *C. elegans* databases. This work was performed in cCAMP (NCBS), Bangalore.

**Microscopy.** DIC images were captured using Nikon NiE microscope and NIS AR software (Japan). Stereoscopic images were obtained through a Nikon SMZ25 stereo microscope (Japan).

**Oligos (5'-3').**

Northern:

*let-7*(WT): AAC TAT ACA ACC TAC TAC CTC A

*let-7(n2853)*: AAC TAT ACA ACC TAC TAT CTC A

*mir-84*: TCT ACA ATA TTA CAT ACT ACC TCA

*mir-48*: TCG CAT CTA CTG AGC CTA CCT CA

*mir-241*: TCA TTT CTC GCA CCT ACC TCA

*mir-77*: TGG ACA GCT ATG GCC TGA TGA A

*mir-237*: AGC TGT TCG AGA ATT CTC AGG GA

*lin-4*: TCA CAC TTG AGG TCT CAG GGA

*mir-90*: AGG GGC ATT CAA ACA ACA TAT CA

*mir-124*: TGG CAT TCA CCG CGT GCC TTA

*mir-75*: TGA AGC CGG TTG GTA GCT TTA A

*mir-1*: TAC ATA CTT CTT TAC ATT CCA

*mir-234*: AAG GGT ATT CTC GAG CAA TAA

*mir-79*: AGC TTT GGT AAC CTA GCT TTA T

$tRNA^{Gly}$: GCT TGG AAG GCA TCC ATG CTG ACC ATT

qPCR:

Primary *mir-77*

Forward: CAT TGT TCG TTT CGC TTT CA

Reverse: CCA ATA ACT GAT TCA ACA TTC CAA

Primary *let-7*

Forward: TCC TAG AAC ACA TCT CCC TTT GA

Reverse: CGC AGC TTC GAA GAG TTC TG

*daf-12* mRNA

Forward: GAT CCT CCG ATG AAC GAA AA

Reverse: CTC TTC GGC TTC ACC AGA AC

*lin-41* mRNA

Forward: GGA TTG TTC GAC ACC AAC G

Reverse: ACC ATG ATG TCA AAC TGC TGT C

*act-1* mRNA

Forward: CAC GGT ATC GTC ACC AAC TG

Reverse: GTA CGT CCG GAA GCG TAG AG

Cloning:

*xrn-2* cDNA (for Baculovirus expression)

Forward Primer: CAA GAA TTC TAA TGG GAG TTC CCG CAT TCT TCA GAT G

Reverse primer: GCG AAG CTT TTA TCT CCA TGA TGA ATT TCC GTG ATA G

*paxt-1* cDNA (for bacterial expression)

Forward Primer: CAT AGG ATC CAT GGG GAA GTT AGA AGA CG

Reverse primer: CGT TCT CGA GTT ATT GGA ATG CAA GAG AGC

*nol-58* cDNA (for bacterial expression)

Forward Primer: CCG GAA TTC ATG CTG GTC CTT TTC GAG G

Reverse primer: AAT GCG GCC GCC TAT TCT TCA TCA GAA TCA GTA TCG G

*miRNasome-1.4* cDNA (for bacterial expression)

Forward Primer: CTG CGG CCG CTA ATG GAA ACC GAA TTT G

Reverse primer: CTG CGG CCG CTC ATT GGT CCT CTG AAA C

Site Directed Mutagenesis:

*xrn-2* cDNA (XRN-2; D234-A, D236-A)

Forward Primer: GCC TCT GCG GAG CCG CCG CCG CCC TTA TTA TGC TCG

Reverse primer: CCG AGC ATA ATA AGG GCG GCG GCG GCT CCG CAG AGG C

Preparation of templates for *in vitro* transcription:

Mature *mir-84* cassette:

Forward primer (T7 Promoter, HH Ribozyme, First 12 *mir-84* nt.)
G TAA TAC GAC TCA CTA TAG GG AGA CAT ACT ACC TCA CTG ATG AGT CCG TGA GGA CGA AAC GGT ACC CGG TAC CGT CTG AGG TAG TAT G

Reverse primer (Mature *mir-84* complementary sequence, 12 nt. complementary region to HH Ribozyme)
TCT ACA ATA TTA CAT ACT ACC TCA GAC GGT ACC GGG

Mature *let-7* cassette:

Forward primer (T7 Promoter, HH Ribozyme, First 12 *let-7* nt.)
G TAA TAC GAC TCA CTA TAG GGAGA CTA CTA CCT CAC TGA TGA GTC CGT GAG GAC GAA ACG GTA CCC GGT ACC GTC TGA GGT AGT AGG

Reverse primer (Mature *let-7* complementary sequence, 12 nt. complementary region to HH Ribozyme)
AAC TAT ACA ACC TAC TAC CTC A GAC GGT ACC GGG

Mature *mir-237* cassette:

Forward primer (T7 Promoter, HH Ribozyme, First 13 *miR-237* nt.)
G TAA TAC GAC TCA CTA TAG GGG AGA CGA GAA TTC TCA GGG ACT GAT AGT CCG TGA GGA CGA AAC GGT ACC CGG TAC CGT CTC CCT GAG AAT TCT CGA ACA GCT

Reverse primer (Mature *mir-237*, 13 nt. Complementary region to HH Ribozyme)
AGC TGT TCG AGA ATT CTC AGG GA GAC GGT ACC GGG

#### Synthetic RNAs (5'-3').

pre-*let-7*: UGA GGU AGU AGG UUG UAU AGU UUG GAA UAU UAC CAC CGG UGA ACU AUG CAA UUU UCU ACC UUA CC

*let-7*: UGA GGU AGU AGG UUG UAU AGU U

*mir-84*: UGA GGU AGU AUG UAA UAU UGU AGA

*lin-4*: UCC CUG AGA CCU CAA GUG UGA

40 nt. RNA: UGA GGU AGU AGG UUG UAU AGU UUA CAA AGU AUU UGA AAA G

15 nt. RNA: UGA GGU AGU AGG UUG

**Supplementary Text 1**

**Developmental consequences of *nol-58*(*RNAi*) in *let-7*(*n2853*) and *N2* worms.**

Most of the *let-7*(*n2853*); *nol-58(RNAi)* animals displayed defects in vulval morphogenesis (75% $\pm$ 2.0% [n=200]), i.e., the vulva did not close during the late L4/young adult stage (**fig. S5A middle panel**). Vulval formation appeared to be delayed for those worms, rather than aborted, as 74% ($\pm$ 2.5% [n=200]) of the *let-7*(*n2853*)*; nol-58*(*RNAi*) adult animals ultimately developed fully closed but protruded vulva (**fig. S5B top** and **middle panels**).

Depletion of NOL-58 in wild type *N2* worms resulted in a plethora of developmental defects like slow growth, larval arrest, delayed vulval morphogenesis, protruded vulva, sterility, arrest in embryonic development etc (**fig. S6A**), and it also led to the formation of precocious alae in ~70% ($\pm$ 4% [n=50]) of the worms (**fig. S6B right panels**), which was again suggestive of higher *let-7* activity.

*N2* Worms undergoing RNAi from L1 stage arrested at L4 (98% $\pm$ 2.0%, [n=200]), and 20% ($\pm$ 2.0%, n=200) showed protruded vulva. When RNAi was initiated at L2 stage, 99% ($\pm$ 1.0%, n=200) of the worms turned into sterile adults with no embryo formation, and 85% ($\pm$ 3.0%, n=200) showed delayed vulval closure, whereas 42% ($\pm$ 3.0%, n=200) of them showed protruded vulva. 95% ($\pm$ 4.0%, n=200) of the worms turned into adults with arrested embryos, when RNAi was initiated at L3 stage. 61% ($\pm$ 3.0%, n=200) of them also showed protruded vulva. Notably, worms undergoing RNAi from L4 stage produced F1 progeny, which did not develop beyond L2 stage and arrested either as L1 (82% $\pm$ 2.0%, [n=200]) or as L2 (18% $\pm$ 1.5%, [n=200]) stage worms.

**Supplementary Text 2**

***nol-58* is a genetic enhancer of *xrn-2***

We used a *xrn-2ts* strain, which shows the phenotypes of a worm undergoing *xrn-2*(*RNAi*), when grown at the non-permissive temperature of 25°C, but grows normally at the permissive temperature (20°C, reference *27*). When L2 staged wild type *N2* worms were grown on the *nol-58*(*RNAi*) plates at 20°C, they showed a number of developmental defects as described before (**Supplementary Text 1**, **fig. S6A**), but attained adulthood (**fig. S7 top right panel**). But the *xrn-2ts* worms growing on *nol-58*(*RNAi*) plates at the permissive temperature of 20°C got arrested at late L3 (75% $\pm$ 3.0% [n=200]) and L4 (23% $\pm$ 1.5% [n=200]) stages, also showed molting defects (43% $\pm$ 5.0% [n=200]; **fig. S7 bottom right panel**), and other anomalies which are characteristic of a *xrn-2ts* worm at the non-permissive temperature (*6, 27*), as well as wild type worm undergoing *xrn-2*(*RNAi*) (reference *5*). Conversely, the same *xrn-2ts* worms growing at the permissive temperature on control plates attained adulthood and showed no defects (**fig. S7 bottom left panel**). These results clearly indicated that *nol-58* genetically interacts with *xrn-2*.

**Supplementary Text 3**

**Micrococcal nuclease efficiently removes all the endogenous RNA from the worm lysate, and after its inactivation allows the detection of only exogenous RNA.**

We performed the pre-*let-7* processing assay as described in **Fig. 2E top panel**, but employed unlabeled pre-*let-7* and subjected the reaction to northern probing using anti-*let-7* probe (**Fig. 2E bottom panel**). Here, in order to eliminate endogenous RNAs in the worm lysates, we treated them with micrococcal nuclease, which would ensure that the northern-detected mature miRNA can only be a derivative of the exogenous pre-miRNA. The same samples were treated with EGTA for chelation of $Ca^{++}$ to terminate the micrococcal nuclease activity, before they were used for the assays. Omission of EGTA failed to detect any band through northern probing (**Fig. 2E bottom panel, lane 5**), which further confirmed efficient removal of endogenous RNAs by the micrococcal nuclease (in **lanes 3, 4 of Fig. 2E bottom panel**).

#### Supplementary Figures

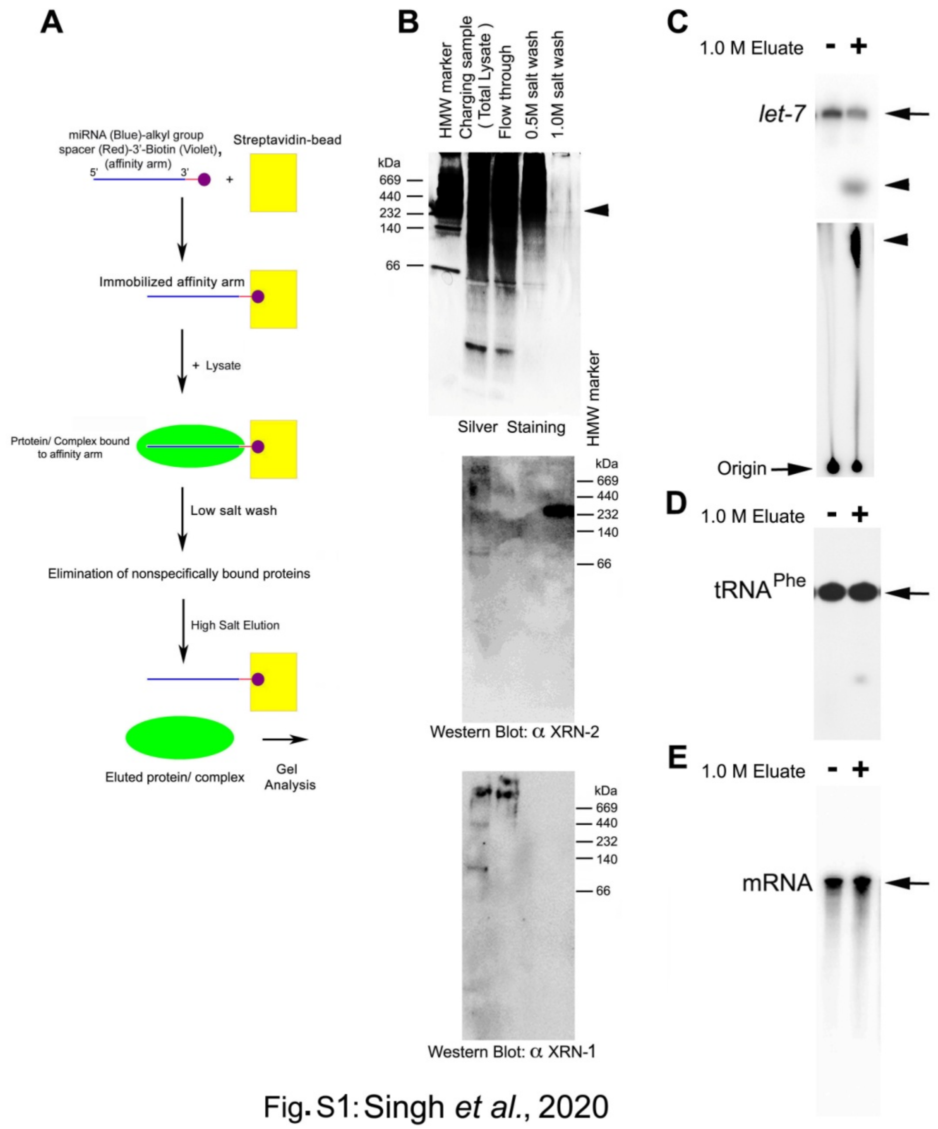


**Fig. S1. RNA affinity purification of a XRN-2 containing complex**

(**A**) Flow diagram for RNA affinity purification, where a phosphorothioate stabilized and appropriately modified 5’-monophosphorylated *let-7* miRNA, as indicated, has been used as an affinity arm.

(**B**) High salt elution fraction (from the RNA affinity chromatography) enriched with a XRN-2 containing high molecular weight band (~260 kDa, middle panel), detected through western blotting. XRN-1 couldn't be detected in the same 1.0 M eluate (bottom panel). Silver staining detected a modest band (top panel, arrowhead) corresponding to the band detected through anti-XRN-2 western blotting (middle panel). Note that anti-XRN-1/2 western blotting detected multiple high molecular weight bands in the total lysate, which were larger than their individual sizes.

**(C)** Incubation of a 5'-radiolabeled *let-7* miRNA (arrow) with the 1.0 M eluate of the RNA affinity chromatography resulted in the formation of a single product on a Urea PAGE (top panel), which was further confirmed as monoribonucleotide (Uridine 5'-monophosphate [UMP], arrowhead, bottom panel) through TLC analysis. The same protein fraction showed very little or insignificant activity on a 5'-radiolabeled tRNA (**D**), and a body-labeled *Renilla* luciferase mRNA (**E**).

Substrate RNAs have been indicated with an arrows, and products with arrowheads.

Representative images from three repeats have been presented.

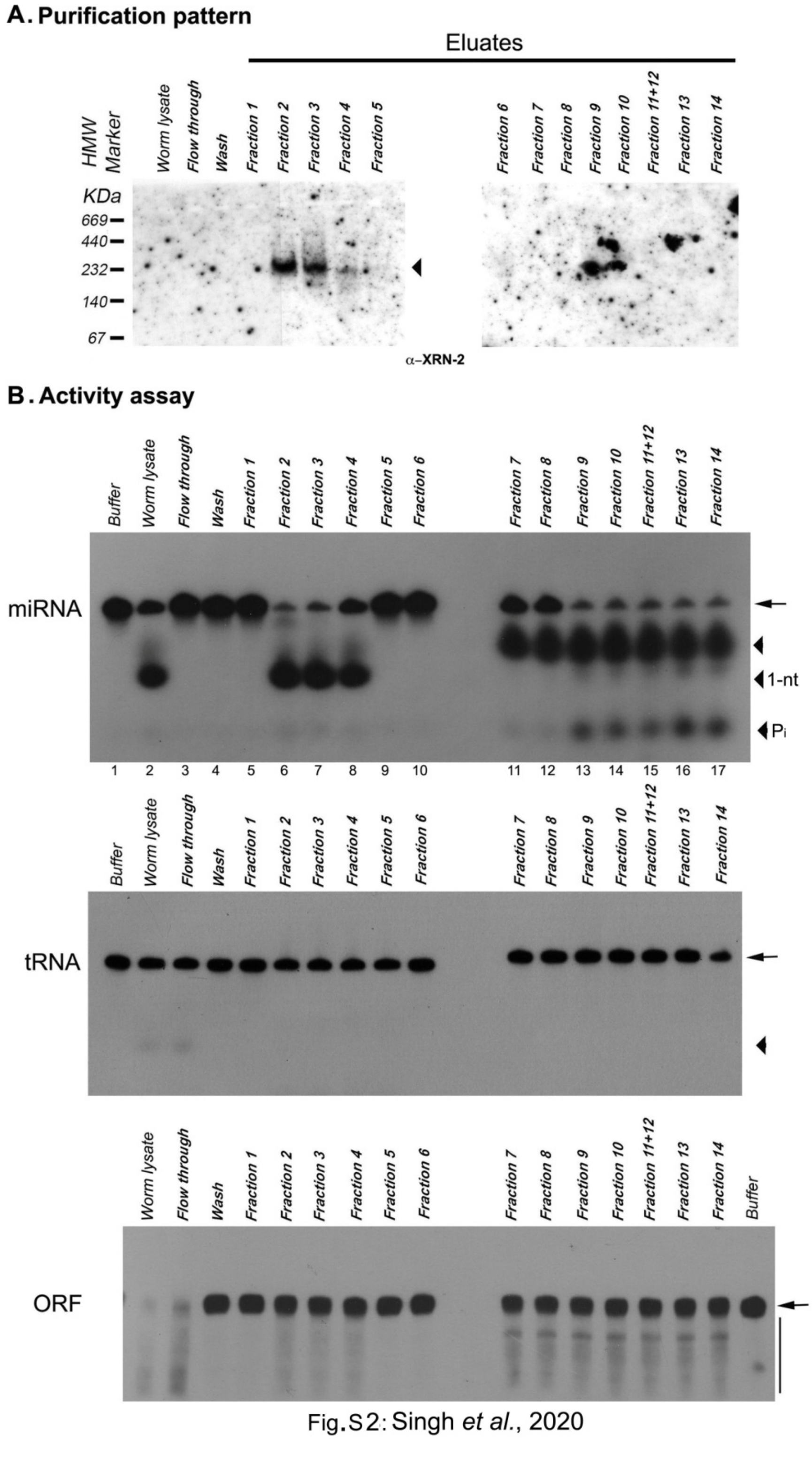


Fig.S2: Singh *et al.*, 2020

**Fig. S2. Fractionation of XRN-2 mediated miRNA turnover activity from total worm lysate**

(**A**) Fractionation of total worm lysate using anion exchange (Capto Q) chromatography. Western blotting detected one XRN-2 containing high molecular weight band (~260 kDa) in fractions 2-4.

(**B**) Activities of the anion exchange chromatography fractions on different RNA substrates, as indicated on the left of each panel. 3'-pCp-labeled and blocked, and 5'-monophosphorylated mature *let-7* substrate (arrow) got efficiently converted into product (Top panel, middle arrowhead), as it did with the worm lysate (compare lanes 2, 6-8). 5'-radiolabeled tRNA (arrow) remained largely unaffected (middle panel), and body-radiolabeled transcript (arrow, bottom panel) of the *Renilla* luciferase open reading frame (ORF, ~1.1 kb) showed a completely different pattern of products (vertical bar, resolved on a 7M urea-3.5% acrylamide gel).

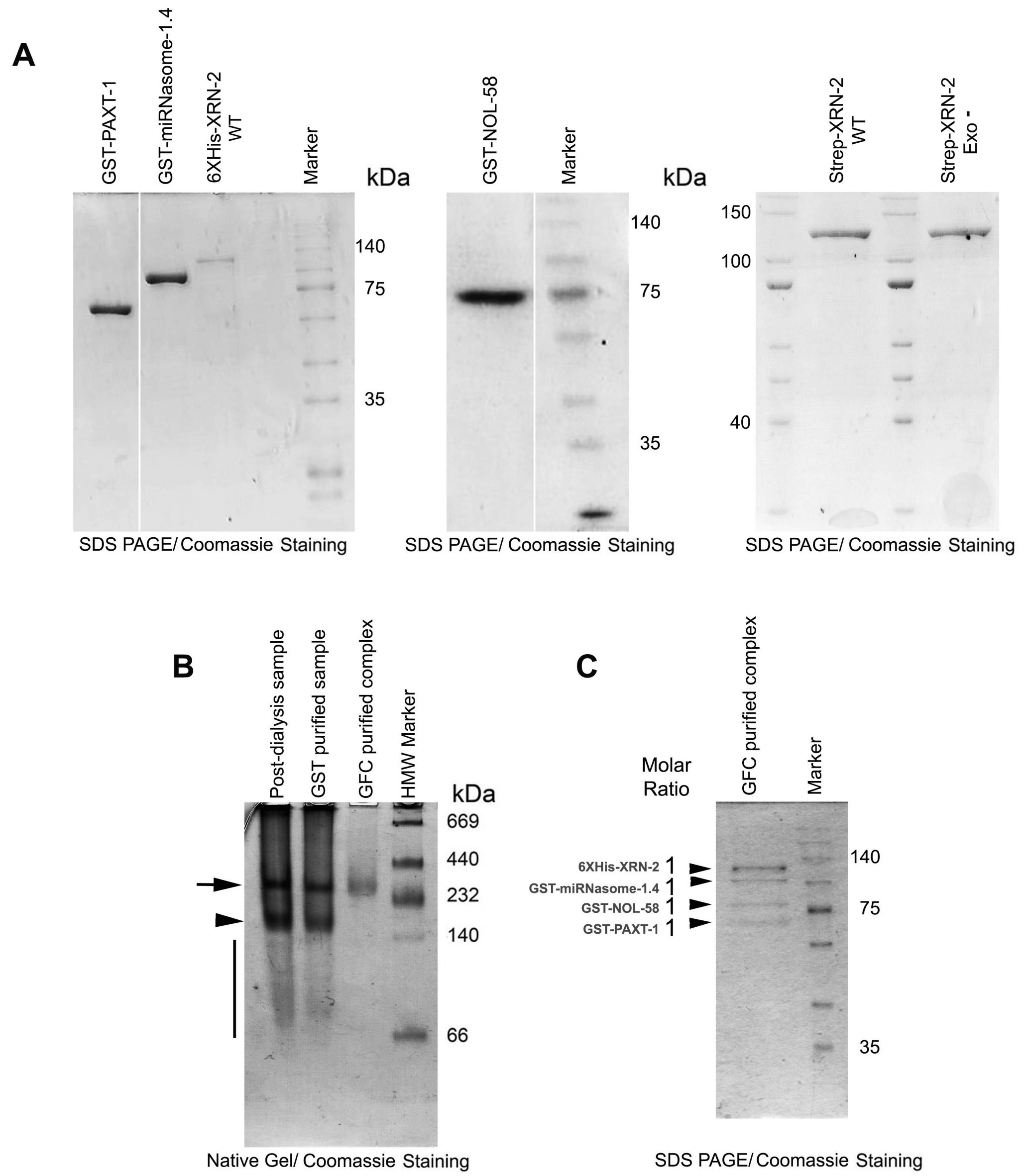


Fig. S3: Singh *et al.*, 2020

**Fig. S3. *in vitro* reconstitution and purification of miRNasome-1**

(**A**) Purified recombinant GST-tagged PAXT-1, NOL-58, and miRNasome-1.4 expressed in *E. coli*, and 6XHis/ Strep tagged XRN-2 (wild type and exoribonuclease mutant) expressed in insect cells, used for *in vitro* reconstitution and *in vitro* assays described elsewhere. Unrelated lanes between GST-PAXT-1 and GST-miRNasome-1.4, GST-NOL-58 and marker were removed.

(**B**) Purified individual recombinant proteins were mixed in equimolar ratio and dialysed overnight, then subjected to affinity purification using Glutathione Sepharose™ 4B resin, followed by Gel Filtration Chromatography (GFC) to separate the *bona fide* complex from sub-complexes and individual proteins. Samples from the aforementioned steps were resolved on a 5% native gel and stained with Coomassie Brilliant Blue. Reconstituted miRNasome-1 is indicated with an arrow, arrowhead indicates to a major sub-complex, and the vertical bar is used to indicate minor sub-complexes and individual component proteins.

**(C)** SDS PAGE analysis of the GFC purified reconstituted miRNasome-1 revealed the presence of four bands in the apparent stoichiometry of 1:1:1:1, with respect to each other. Note that the presence of salt has resulted in slower than expected migration for the proteins.

Representative images from three repeats have been presented.

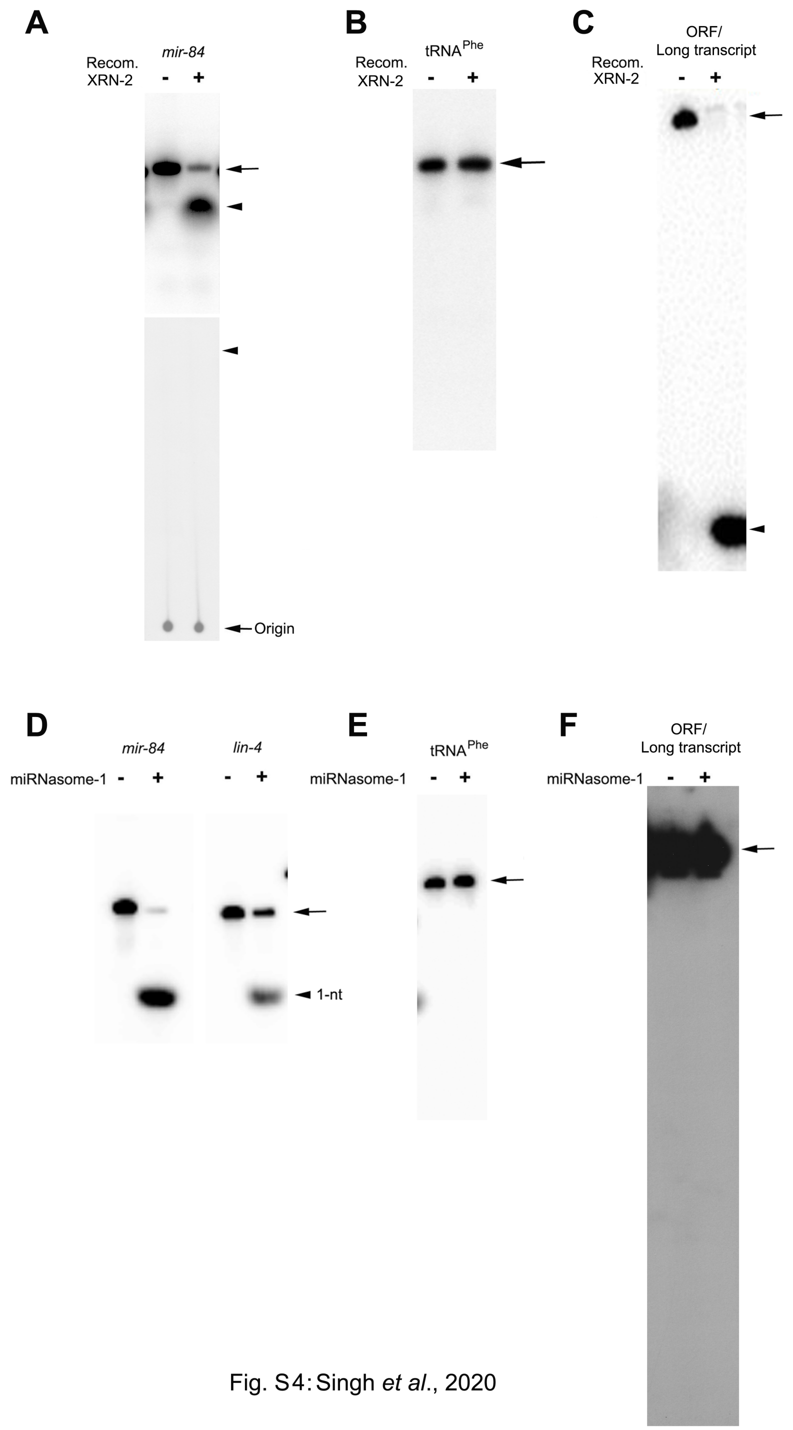


**Fig. S4. Recombinant XRN-2 lacks specificity (A-C), whereas, *in vitro* reconstituted miRNasome-1 shows specific activity (D-F) in *in vitro* turnover assays.**

(**A**) Activity of recombinant XRN-2 (10 ng, ~9.0 nM) on a 5'-radiolabeled mature miRNA (*mir-84*) yields a product (arrowhead, **top panel**), which doesn't form a spot depicting monoribonucleotides on a TLC plate (**bottom panel**). An arrowhead indicates the expected level at which monoribonucleotides should migrate on the TLC plate.

(**B**) Recombinant XRN-2 (~9.0 nM) shows very little or no activity on a 5'-radiolabeled tRNA (arrow).

(**C**) Upon incubation with recombinant XRN-2 (~9.0 nM), a body-radiolabeled *Renilla* luciferase ORF (~1100-nt) gets converted into product(s), indicated with an arrowhead, which migrate ahead of the dye-front of the gel (3.5% Urea-PAGE).

The substrate RNA concentration in all the above reactions was 10 nM.

(**D**) Activity of purified reconstituted miRNasome-1 (100 ng, ~40 nM) on different 5'-radiolabeled mature miRNAs (arrow), as indicated. It is to be noted that product (arrowhead) formation was modest for *lin-4*, which is consistent with the activity of the endogenous miRNasome-1 (**Fig. 1F**).

(**E**) Purified reconstituted miRNasome-1 (~40 nM) shows no activity on a 5'-radiolabeled tRNA, and on a body-radiolabeled *Renilla* luciferase ORF (**F**).

Substrate RNA concentration in all the above reactions with miRNasome-1 was in five fold molar excess, and substrates are indicated with arrows, whereas products with arrowheads.

Representative images from three repeats have been presented.

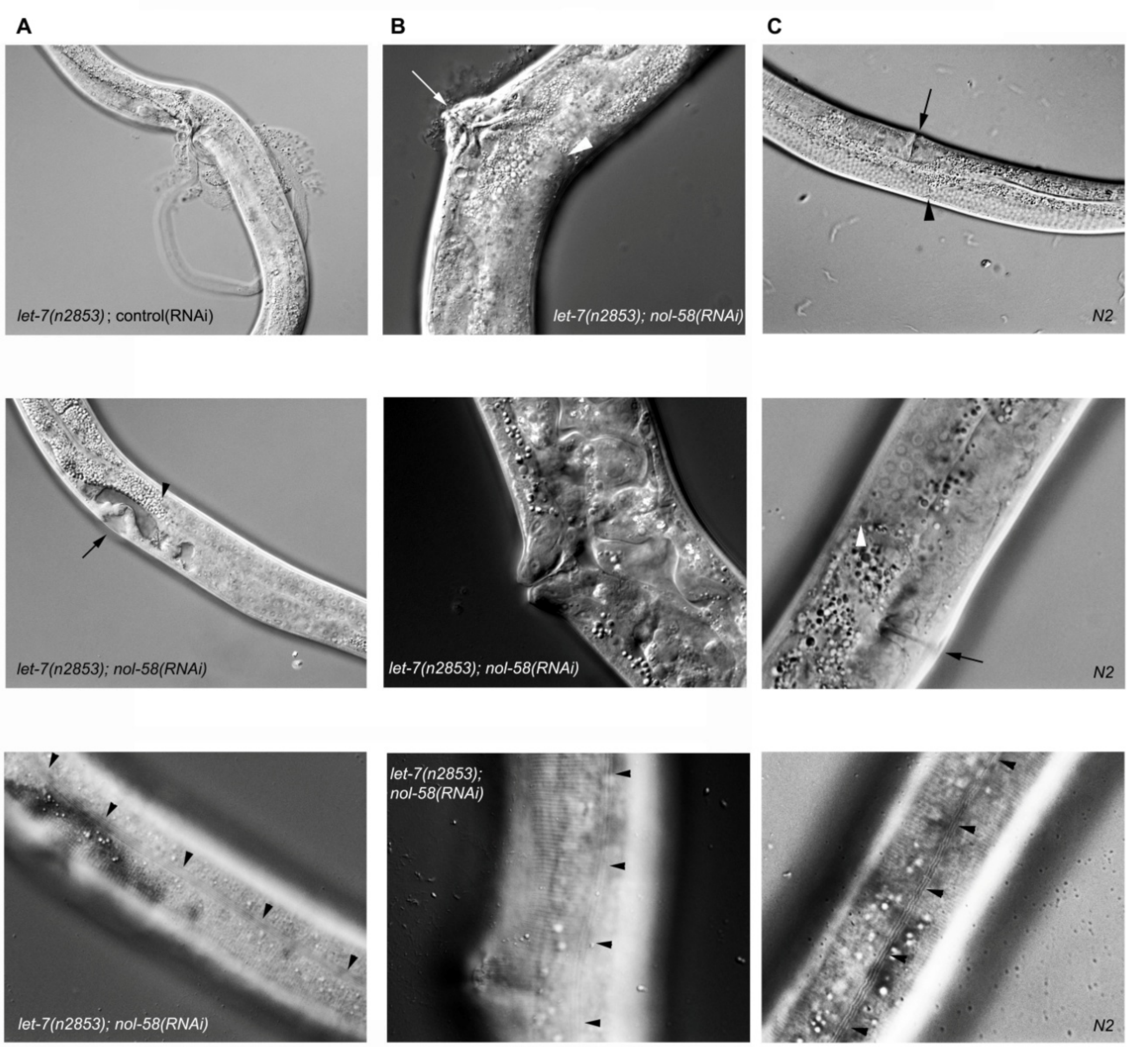


**Fig. S5. *nol-58*(*RNAi*) suppresses the bursting phenotype of *let-7(n2853)* worms**

(**A**) Synchronized L1 larvae of *let-7(n2853)* worms were grown on OP50 plates at 25°C until they became late L1, and then they were transferred either on empty vector RNAi plates (control) or *nol-58(RNAi)* plates and grown at 25°C to young adult stage. Worms growing on control plates (**top panel**, at 37-39 hrs) died by bursting through their vulvae, whereas *let-7(n2853); nol-58(RNAi)* animals did not (**middle panel**, at ~41 hrs). Arrow points to the vulva, arrowhead to distal gonadal tip, indicating correct late L4 stage of an animal with delayed vulval closure. Continuous alae were visible in the experimental animals with delayed vulval closure, even at late L4 stage (indicated by arrowheads, **bottom panel**)

(**B**) **Top panel** shows a *let-7(n2853); nol-58(RNAi)* worm, at 50 hrs, with closed, protruded, but intact vulva (arrow). The distal tip of one gonadal arm is pointed by an arrowhead, which indicates to the correct adult stage of the animal. **Middle panel** depicts the closed, protruded, but intact vulval architecture at 50 hrs. **Bottom panel** shows development of continuous alae (indicated by arrowheads) in an adult worm with protruded vulva.

(**C**) A wild type *N2* worm with closed vulva (arrow, **top panel**) at young adult stage, indicated by the 'just touching each other' position of the distal tips of the two gonadal arms (arrowhead). **Middle panel** shows a worm at late L4 stage, but already with a closed vulva (arrow). Arrowhead points to distal gonadal tip, indicating correct late L4 stage. **Bottom panel** depicts the development of continuous alae in the same worm shown in the middle panel.

Representative images from three or more repeats have been presented.

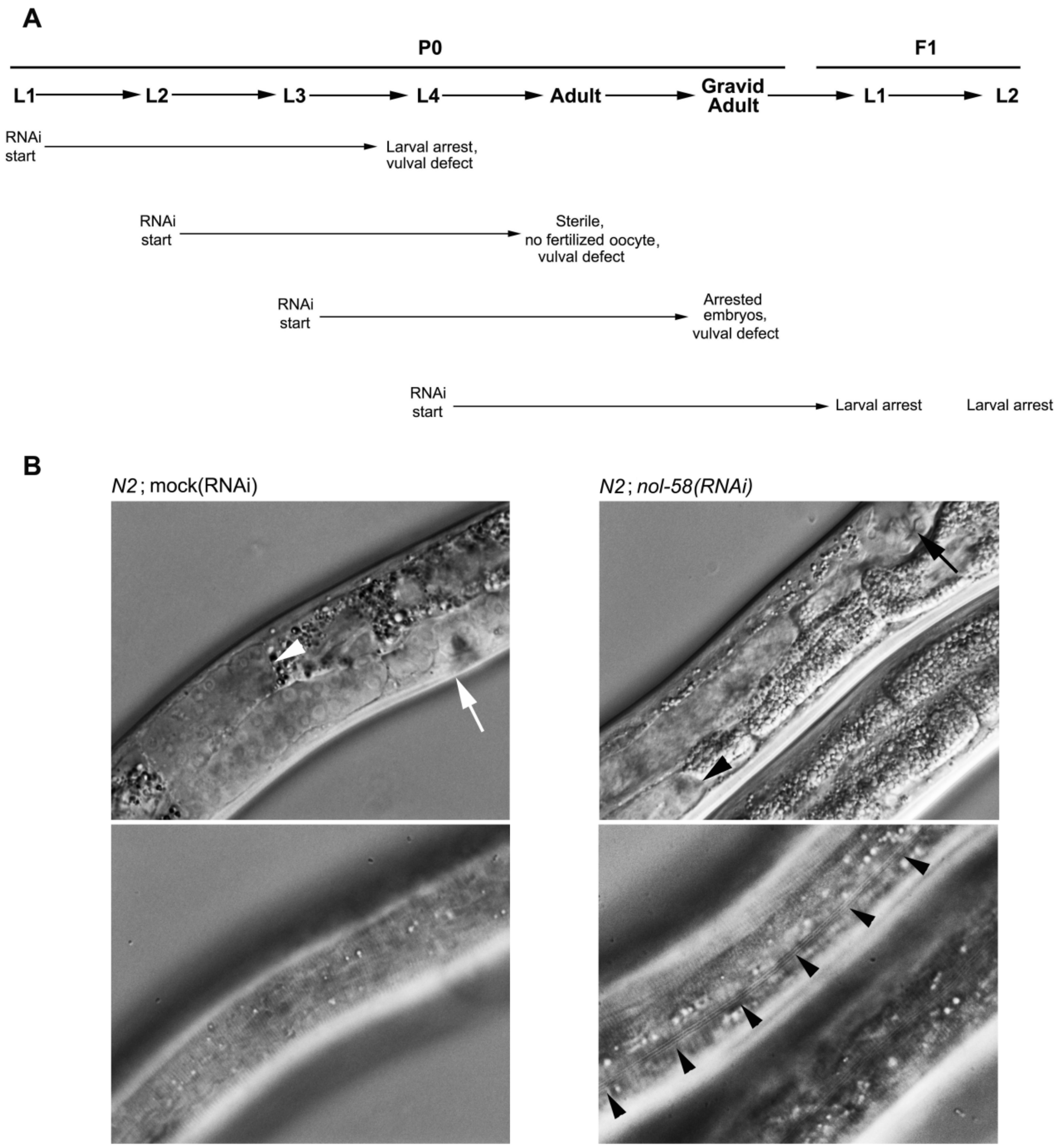


Fig. S6 : Singh *et al.*, 2020

**Fig. S6. Developmental consequences of *nol-58*(*RNAi*) on wild type *N2* worms**

**(A)** Effects on the worms exposed to RNAi plates from a given larval stage, as indicated (RNAi start). The effects are listed below the life cycle stages where they were observed. Detailed description and statistics have been furnished in the supplementary text 1.

**(B)** Precocious alae formation (indicated through arrowheads, **right bottom panel**) in a worm undergoing depletion of NOL-58. The worm is at early-mid L4 stage as depicted by the gonadal arm position (arrowhead, **right top panel**). An arrow indicates to the open vulva. Alae is not visible in a control worm at mid L4 stage (**left bottom panel**), where gonadal arm position (**left top panel**) confirmed correct staging. The distal end of the gonadal arm is indicated by an arrowhead, and the vulva by an arrow. Representative images from three or more repeats have been presented.

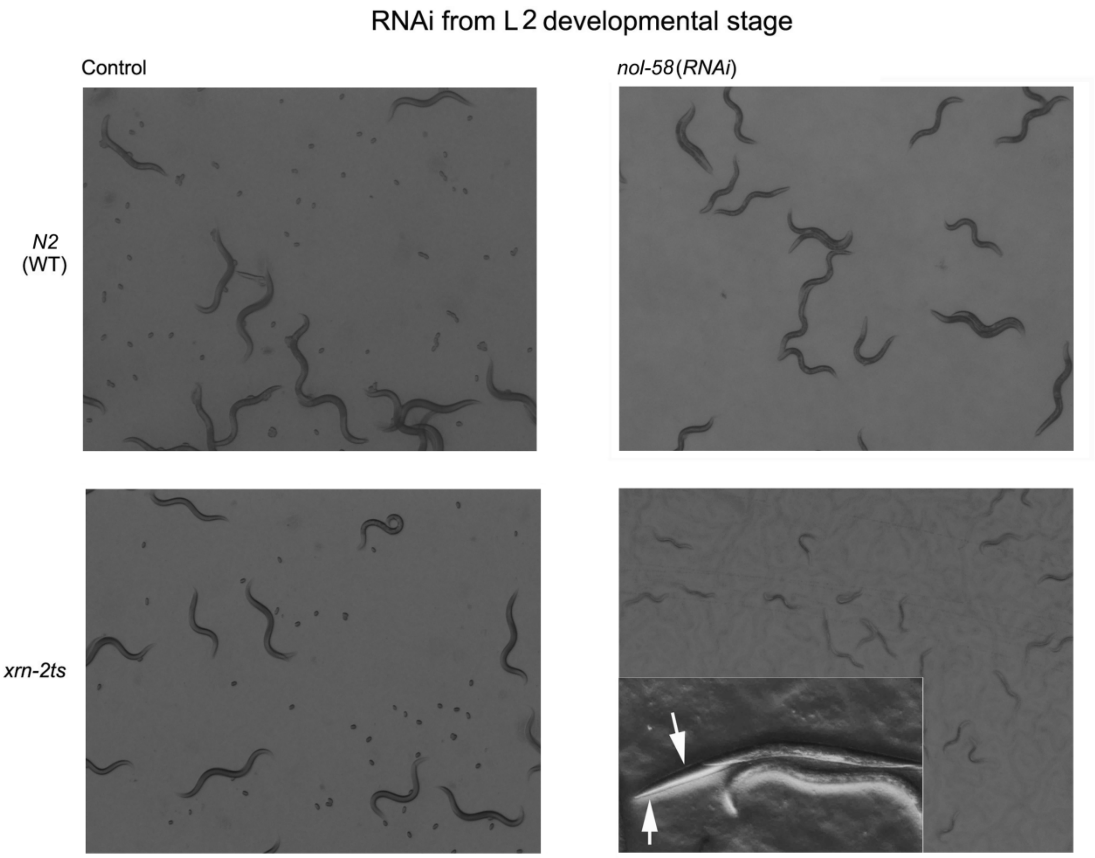


Fig. S7: Singh *et al.*, 2020

**Fig. S7. *nol-58* is a genetic enhancer of *xrn-2***

Wild type *N2* or *xrn-2ts* worms were grown on control or *nol-58*(*RNAi*) plates, as indicated, from L2 stage at 20°C. Wild type worms (**top left panel**) on control plates turned gravid adult and laid embryos, whereas the ones on *nol-58*(*RNAi*) plates were largely grown into sterile adults (**top right panel**). *xrn-2ts* worms growing on control plates turned gravid and laid embryos (**bottom left panel**), albeit they had grown little slower than the wild type worms. Whereas, *xrn-2ts* worms growing on *nol-58*(*RNAi*) plates got arrested in late L3/ early L4 stages (**bottom right panel**), and 43% (+5%, n=200) of the worms showed molting defect, which has also been observed for wild type worms undergoing *xrn-2*(*RNAi*) (data not shown, and references *5, 6*). An enlarged picture of a L3/L4 worm with unshed cuticle (arrows) is shown in the inset. Representative images from three or more repeats have been presented.

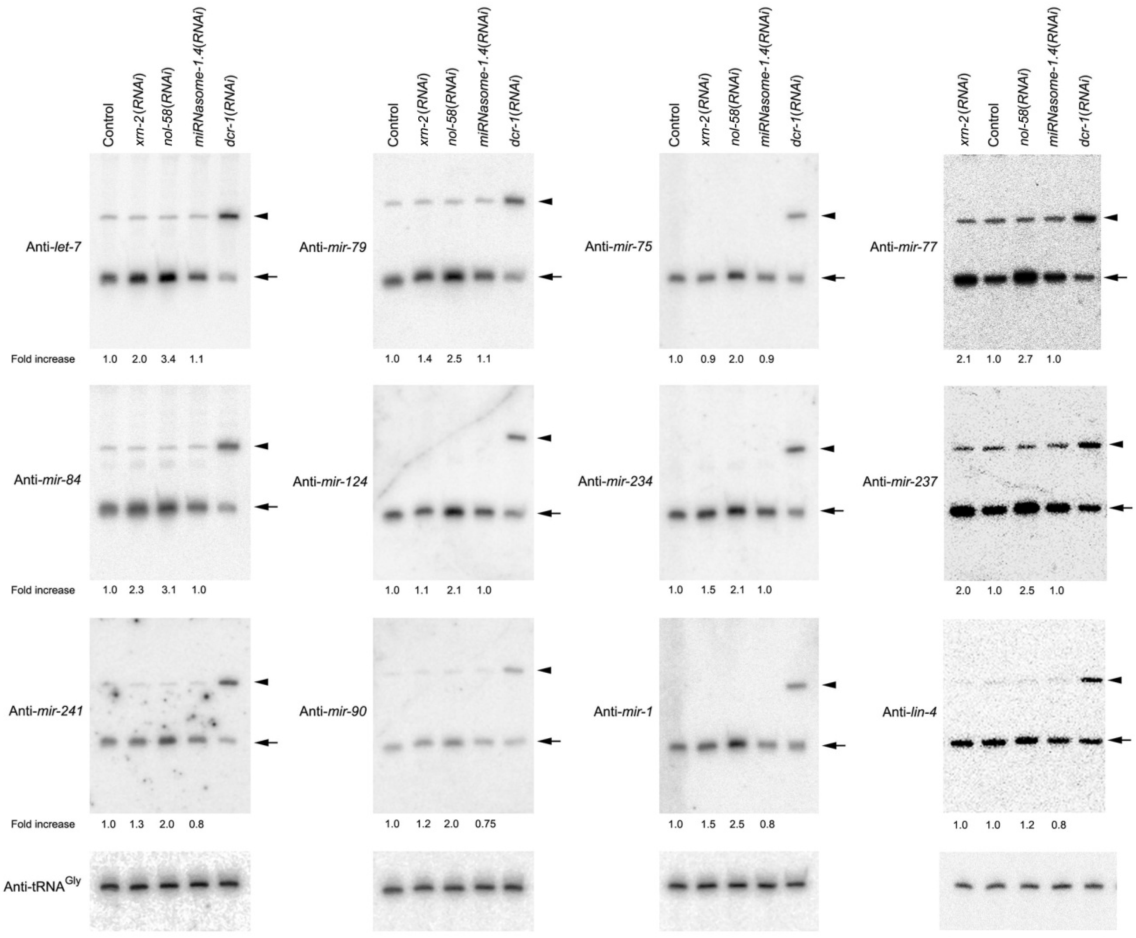


Fig. S8: Singh *et al.*, 2020

**Fig. S8. Depletion of miRNasome-1 components affects only mature miRNA levels *in vivo*.** Northern blotting of total RNA extracted from wild type *N2* worms exposed to different RNAi, as indicated. *nol-58*(*RNAi*), as well as *xrn-2*(*RNAi*), leads to accumulation of mature miRNAs (arrow) to different extents. The normalized fold increase values, relative to the control, are provided below the respective lanes, where tRNA$^{Gly}$ was used as the normalizing control. Note that pre-miRNAs (arrowhead), wherever they could be detected, remained by and large unchanged, except in the *dcr-1*(*RNAi*) samples, where pre-miRNAs accumulated heavily with concomitant depletion in the levels of mature miRNAs. Four different blots were used, which were stripped and re-probed. Representative images from three or more repeats have been presented.

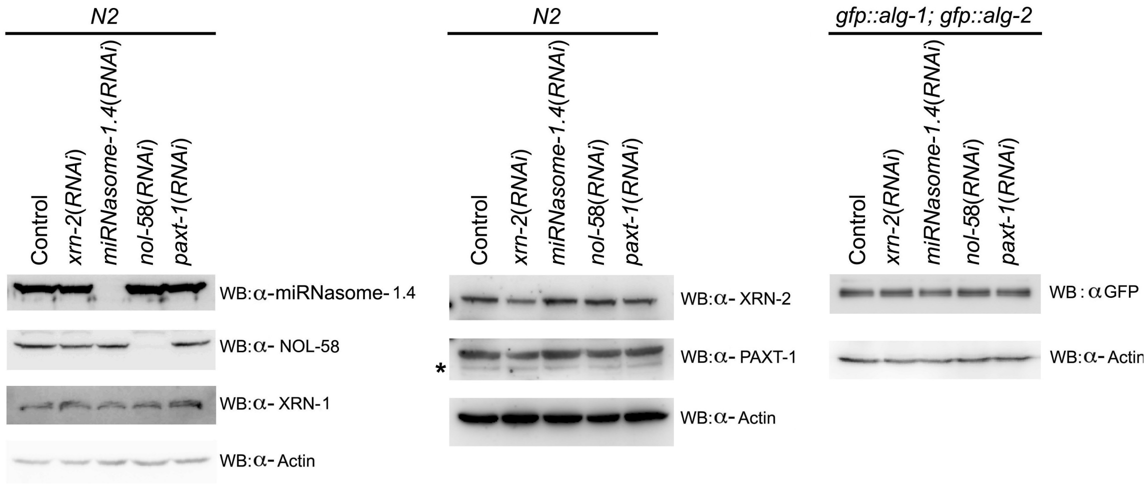


Fig. S9: Singh *et al.*, 2020

**Fig. S9. RNAi depletion of miRNasome-1 components and their effects on other relevant proteins.** Western blotting was performed using lysates made from worms (*N2* or *gfp::alg-1; gfp::alg-2*) undergoing mock (control) or specific RNAi, as indicated. Depletion of NOL-58 and miRNasome-1.4 didn't affect other candidates, as indicated (**left and middle panel**). Under the employed RNAi conditions, PAXT-1 showed modest depletion (**middle panel**). Asterisk indicates to a cross-reacting protein detected by anti-PAXT-1 antibody. XRN-1 (**left panel**), and ALG-1; ALG-2 (**right panel**) remained unaffected upon depletion of miRNasome-1 subunits. Actin served as loading control. Representative images from three or more repeats have been presented.

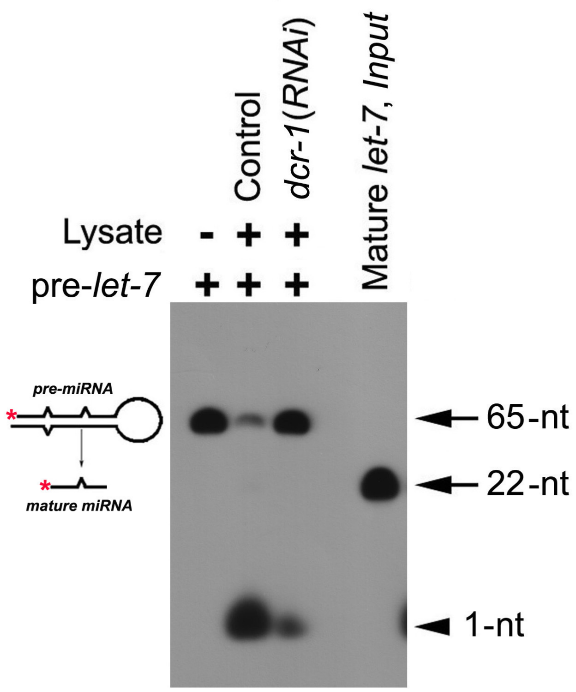


Fig. S10: Singh *et al*., 2020

**Fig. S10. Dicer-dependent product formation in a pre-*let-7* processing assay.** RNAi depletion of DCR-1 inhibits product (1-nt) formation and stabilizes the substrate pre-*let-7* (65-nt) in a pre-*miRNA* processing assay. The 5'-PNK-radiolabeled (red asterisk) pre-*miRNA* and the *in vitro* diced/ processed mature miRNA from the precursor are depicted as a cartoon on the left. A representative image from three repeats has been presented.

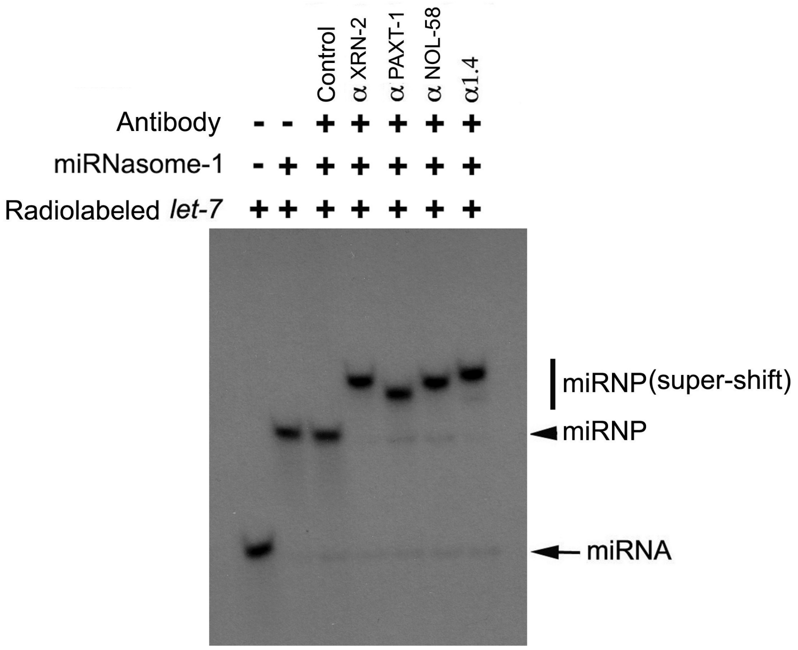


Fig. S11: Singh *et al*., 2020

**Fig. S11. Radiolabeled *let-7* shows super-shift upon incubation with miRNasome-1, and the antibodies against its subunits in an electrophoretic mobility shift assay.** 5'-radiolabeled PTO stabilized synthetic *let-7* (10 fmol, 1.0 nM) forms a distinct miRNP (indicated by arrowhead) upon incubation with purified miRNasome-1 (250 ng, ~100 nM). The *let-7*-miRNasome-1 miRNP gets super-shifted upon further incubation with specific antibodies (0.5 µg, ~350 nM), as indicated. A representative image from three repeats has been presented.

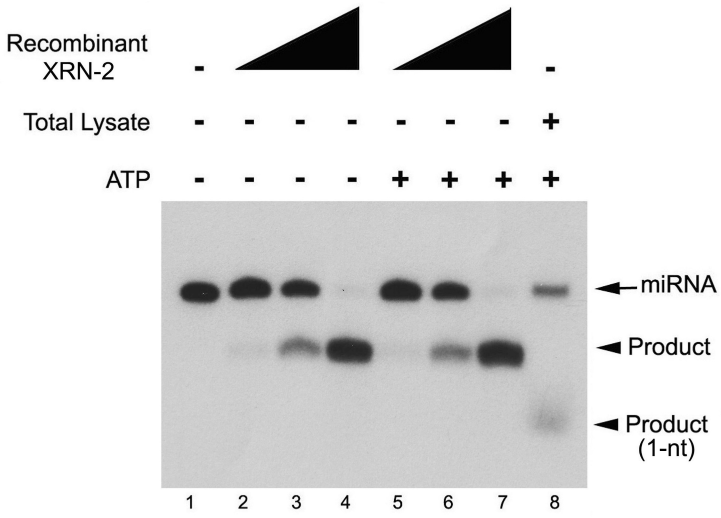


Fig. S12: Singh *et al.*, 2020

**Fig. S12. Mutation of the exoribonuclease active site residues (exo$^-$; D234-A, D236-A) abolishes monoribonucleotide formation by XRN-2, but endoribonucleolytic product formation remains unchanged**. 10 nM body-radiolabeled *mir-84* was subjected to *in vitro* miRNA turnover assay performed with purified exo$^-$ recombinant XRN-2 (in an ascending concentration of 0.9 nM to 90 nM) or total lysate in the absence or presence of ATP, as indicated. The lower arrowhead indicates to monoribonucleotides (lane 8). The reactions with the recombinant proteins were supplemented with 100 nM cold substrate miRNA (*mir-84*), which facilitates monoribonucleotide formation. A representative image from three repeats has been presented.

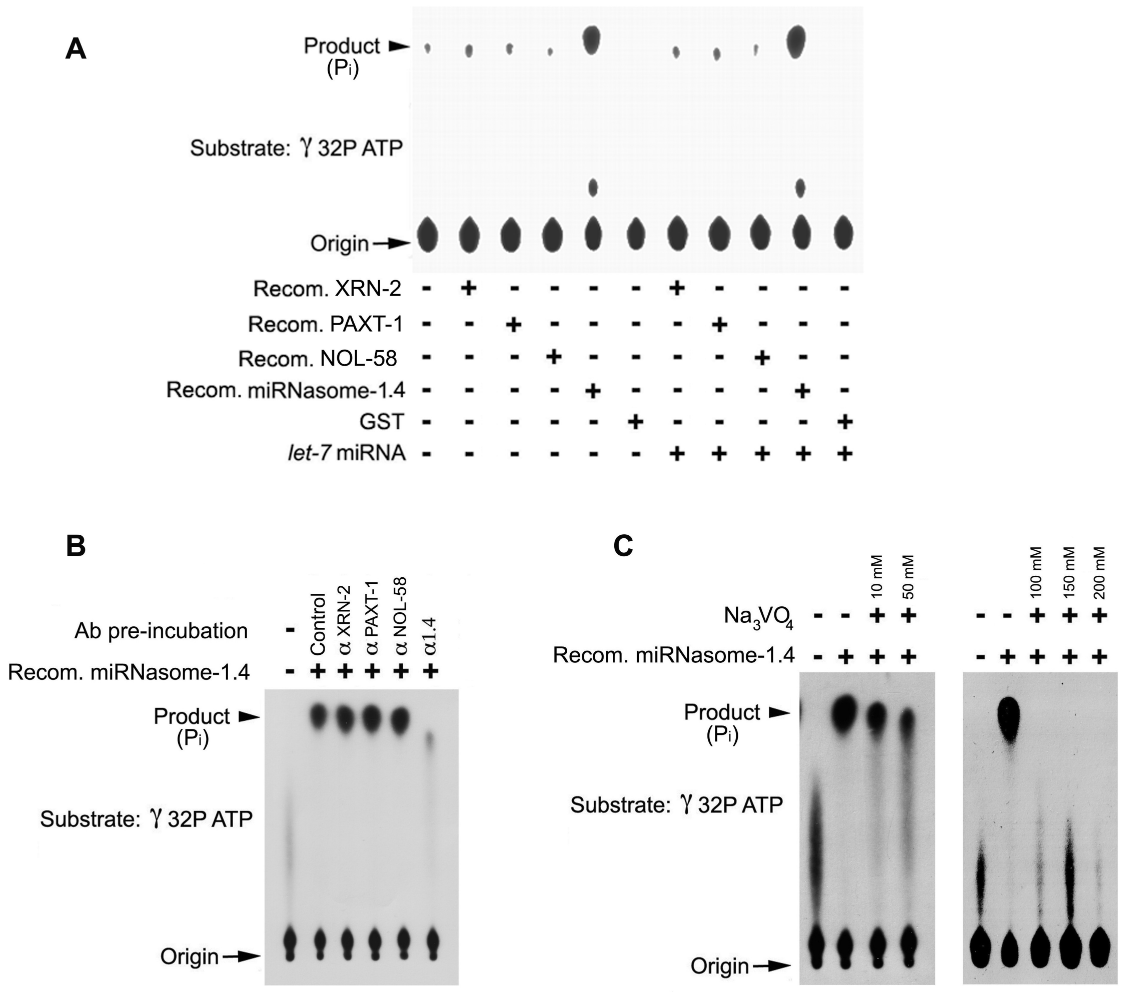


Fig. S13 : Singh *et al*, 2020

**Fig. S13. Specific inhibition of the ATP hydolysing activity of miRNasome-1.4**

**(A)** Recombinant miRNasome-1.4, and no other recombinant versions of the miRNasome-1 subunits, shows ATP hydrolysing activity, and the activity remains unchanged in the presence of a miRNA. **(B)** ATP hydrolysing activity of recombinant miRNasome-1.4 protein gets heavily diminished upon pre-incubation of the protein with anti-miRNasome-1.4 antibody only. (**C**) Sodium orthovanadate inhibits the ATP hydrolysing activity of miRNasome-1.4. Representative images from three repeats have been presented.

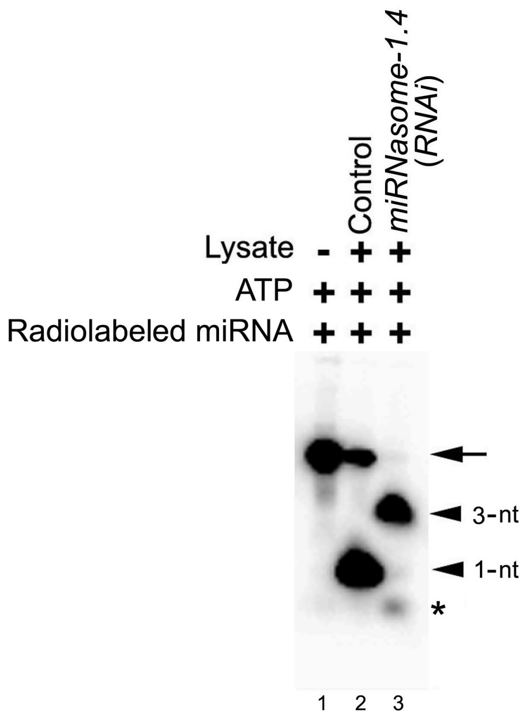


Fig.S14: Singh *et al*., 2020

**Fig. S14. Depletion of miRNasome-1.4 leads to endoribonucleolytic product formation in a miRNA turnover assay in the presence of ATP.** 5'-radiolabeled *mir-84* was incubated with control and miRNasome-1.4 depleted lysates, as indicated, and the reactions were resolved on a urea PAGE. Knock-down lysate resulted in a >90% depletion in monoribonucleotide formation (bottom arrowhead), alongside the accumulation of an endoribonucleolytic product (top arrowhead), which is absent in the control lysate. Asterisk indicates to a minor accumulation of free phosphate in lane three. A representative image from three repeats has been presented.

**Table S1.** Identification of the protein subunits of miRNasome-1 using mass spectrometry

| Mass spectrometry of miRNasome-1 | | | | | | |
|---|---|---|---|---|---|---|
| Accession | Description | | Protein Score+ | Coverage^ | Unique Peptides# | PSM's* |
| 3873676 | Protein B0024.11 | Replicate 1 | 623.89 | 28.08 | 13 | 14 |
| | | Replicate 2 | 811.12 | 38.99 | 17 | 21 |
| 189309813 | Protein XRN-2 | Replicate 1 | 581.70 | 18.26 | 12 | 14 |
| | | Replicate 2 | 564.45 | 15.38 | 13 | 15 |
| 351051006 | Protein NOL-58 | Replicate 1 | 509.79 | 28.95 | 10 | 11 |
| | | Replicate 2 | 218.99 | 15.81 | 6 | 7 |
| 25004996 | Protein PAXT-1 | Replicate 1 | 126.78 | 14.63 | 3 | 3 |
| | | Replicate 2 | 121.69 | 11.04 | 2 | 2 |

+**Protein Score:** The sum of the ion scores of all peptides that were identified.

^**Coverage:** The percentage of the protein sequence covered by identified peptides.

### **Unique Peptides:** The number of peptide sequences that are unique to a protein group. These are the peptides that are common to the proteins of a protein group, and which do not occur in the proteins of any other group.

* **PSM's:** The number of peptide spectrum matches. The number of PSM's is the total number of identified peptide spectra matched for the protein.

Table S1: Singh *et al.*, 2020
